# Specific genomic targeting of the RNF12/RLIM E3 ubiquitin ligase selectively programmes developmental transcription

**DOI:** 10.1101/2023.06.14.544957

**Authors:** Carmen Espejo-Serrano, Catriona Aitken, Beatrice F. Tan, Danielle G. May, Rachel J. Chrisopulos, Kyle J. Roux, Samuel G. Mackintosh, Joost Gribnau, Francisco Bustos, Cristina Gontan, Greg M. Findlay

## Abstract

The E3 ubiquitin ligase RNF12/RLIM controls developmental gene expression and is mutated in the X-linked intellectual disability disorder Tonne-Kalscheuer syndrome (TOKAS). However, the mechanisms by which RNF12 E3 ubiquitin ligase activity controls specific gene expression signatures are not known. Here, we show that chromatin forms a regulatory platform for RNF12 substrate ubiquitylation and transcriptional patterning. RNF12 is recruited to specific genomic regions via a distinct consensus sequence motif, which enables targeting to key transcription factor substrate REX1. Mechanistically, RNF12 chromatin recruitment is largely REX1 independent, but is achieved via the conserved basic region (BR) adjacent to the RING domain. This region is critical for REX1 ubiquitylation on chromatin and downstream RNF12-dependent gene regulation. Furthermore, we find that RNF12 N-terminal sequences suppress chromatin recruitment and substrate ubiquitylation, uncovering a previously unappreciated autoinhibitory mechanism that governs genome targeting. Taken together, our results provide insight into mechanisms by which selective substrate targeting of an E3 ubiquitin ligase enables specific programming of gene expression.

## Introduction

Protein ubiquitylation is a critical post-translational modification that controls all aspects of biology (Kulathu & Komander, 2012; Oh, Akopian, & Rape, 2018) As a result, E3 ubiquitin ligases, which select substrates for ubiquitylation, serve as regulatory gatekeepers for myriad biological processes, including biologically critical functions such as protein homeostasis and quality control, cell cycle and the DNA damage response (Kulathu & Komander, 2012; Oh et al., 2018). Ubiquitylation also orchestrates signalling events, for example in immune cell signalling (Bhoj & Chen, 2009; Popovic, Vucic, & Dikic, 2014). Therefore, dysregulation of E3 ubiquitin ligases has been implicated in many human diseases, such as cancer, disorders of the immune system and developmental disorders (Ciechanover & Brundin, 2003; Popovic et al., 2014; Rape, 2018).

A key function of protein ubiquitylation is in control of gene expression and cell identity, decision-making processes that frequently go awry in disease. This is exemplified by the E3 ubiquitin ligase RNF12/RLIM, which controls developmental gene expression (Bustos et al., 2020; Segarra-Fas et al., 2022; Zhang et al., 2012) and X-chromosome inactivation (Barakat et al., 2011; Gontan et al., 2012; Gontan et al., 2018; Jonkers et al., 2009; Shin et al., 2010), and is mutated in the X-linked intellectual disability disorder Tonne-Kalscheuer syndrome (TOKAS) (Bustos et al., 2021; Frints et al., 2019; Hu et al., 2016; Tønne et al., 2015). RNF12 variants identified in TOKAS patients disrupt RNF12 E3 ubiquitin ligase activity (Bustos et al., 2018; Frints et al., 2019), suggesting that an RNF12 dependent ubiquitin signalling pathway goes awry to cause intellectual disability in these individuals.

RNF12 regulates highly specific gene expression programmes involved in X-chromosome inactivation (Barakat et al., 2011; Gontan et al., 2012; Gontan et al., 2018; Jonkers et al., 2009; Shin et al., 2010), neurodevelopment (Bustos et al., 2020) and gametogenesis (Segarra-Fas et al., 2022). This occurs largely via ubiquitylation and resulting proteasomal degradation of the transcriptional regulator ZFP42/REX1 (Bustos et al., 2020; Gontan et al., 2012; Segarra-Fas et al., 2022). However, beyond this, the molecular details of how RNF12 controls specific gene expression signatures with exquisite precision and accuracy remain unclear. For example, it is not known whether RNF12 engages REX1 specifically on chromatin and/or at specific sites, such as transcriptionally active promoters. Furthermore, the broader role that chromatin context plays in regulation of RNF12 activity towards REX1 has not been studied.

Here, we show that chromatin forms a platform for RNF12 substrate ubiquitylation and transcriptional patterning. We find using proximity labelling that RNF12 engages chromatin components, and ChIP-seq analyses reveal that RNF12 is recruited to specific chromatin regions. In particular, RNF12 is recruited along with REX1 substrate at specific genomic regions such as target gene promoters, leading to REX1 ubiquitylation and gene regulation. Mechanistically, RNF12 engages both chromatin and REX1 via the conserved basic region adjacent to the RING domain, which is critical for efficient REX1 binding and ubiquitylation, and RNF12-dependent gene regulation. Furthermore, RNF12 N-terminal sequences suppress chromatin recruitment and substrate ubiquitylation, uncovering a previously unappreciated autoinhibitory mechanism. Taken together, our results provide insight into mechanisms by which chromatin targeting of an E3 ligase can coordinate catalytic activity and delivery to substrate, enabling implementation of an exquisitely specific gene expression programme.

## Results

### RNF12/RLIM proximity-induced labelling mass spectrometry identifies REX1 substrate and other chromatin associated components

A key function of RNF12 is ubiquitylation and resulting proteasomal degradation of developmental transcriptional regulators (Gao, Wang, Cai, Zhu, & Yu, 2016; Gontan et al., 2012; Her & Chung, 2009; Ostendorff et al., 2002; Fang Wang et al., 2019; Zhang et al., 2012). Chief among these is the transcription factor ZFP42/REX1 (Gontan et al., 2012; Gontan et al., 2018), which patterns developmental gene expression during embryonic stem cell differentiation and is associated with developmental abnormalities (Bustos et al., 2020; Segarra-Fas et al., 2022). However, the mechanisms by which RNF12 is targeted to substrates such as REX1 to control gene expression remain unclear but are critical to understand RNF12 regulation and function in normal and disease states.

To address this question, we took a proximity ligation approach to identify RNF12 proximal proteins. TurboID labelling (Branon et al., 2018) is a proximity ligation-based method that tethers a promiscuous biotin ligase to a protein of interest to rapidly biotinylate and identify proximal proteins. Thus, we fused TurboID machinery to the RNF12 N-terminus to identify proteins that are specifically labelled by RNF12 proximity. RNF12 TurboID and TurboID machinery alone were inducibly expressed in mouse embryonic stem cells (mESCs) in triplicate, incubated with biotin to induce proximity labelling, and treated with MG132 to stabilise RNF12 proximal proteins that might otherwise be targeted for proteasomal degradation. Correct expression and nuclear localisation of HA-TurboID RNF12 was confirmed by immunoblotting (Fig. 1A) and immunofluorescence (Fig. 1B). RNF12 proximal proteins were then identified by streptavidin pull-down and mass-spectrometry, and peptides and proteins quantified to determine fold-change and statistical significance. Proteins whose labelling is increased >2-fold in RNF12 TurboID samples relative to TurboID controls were pinpointed (285 proteins; Fig. 1C & Table S1); proof of principle for this utility of this approach to identify RNF12 proximal proteins was provided by identification of known substrate REX1/ZFP42 (Fig. 1C).

**Fig. 1.**
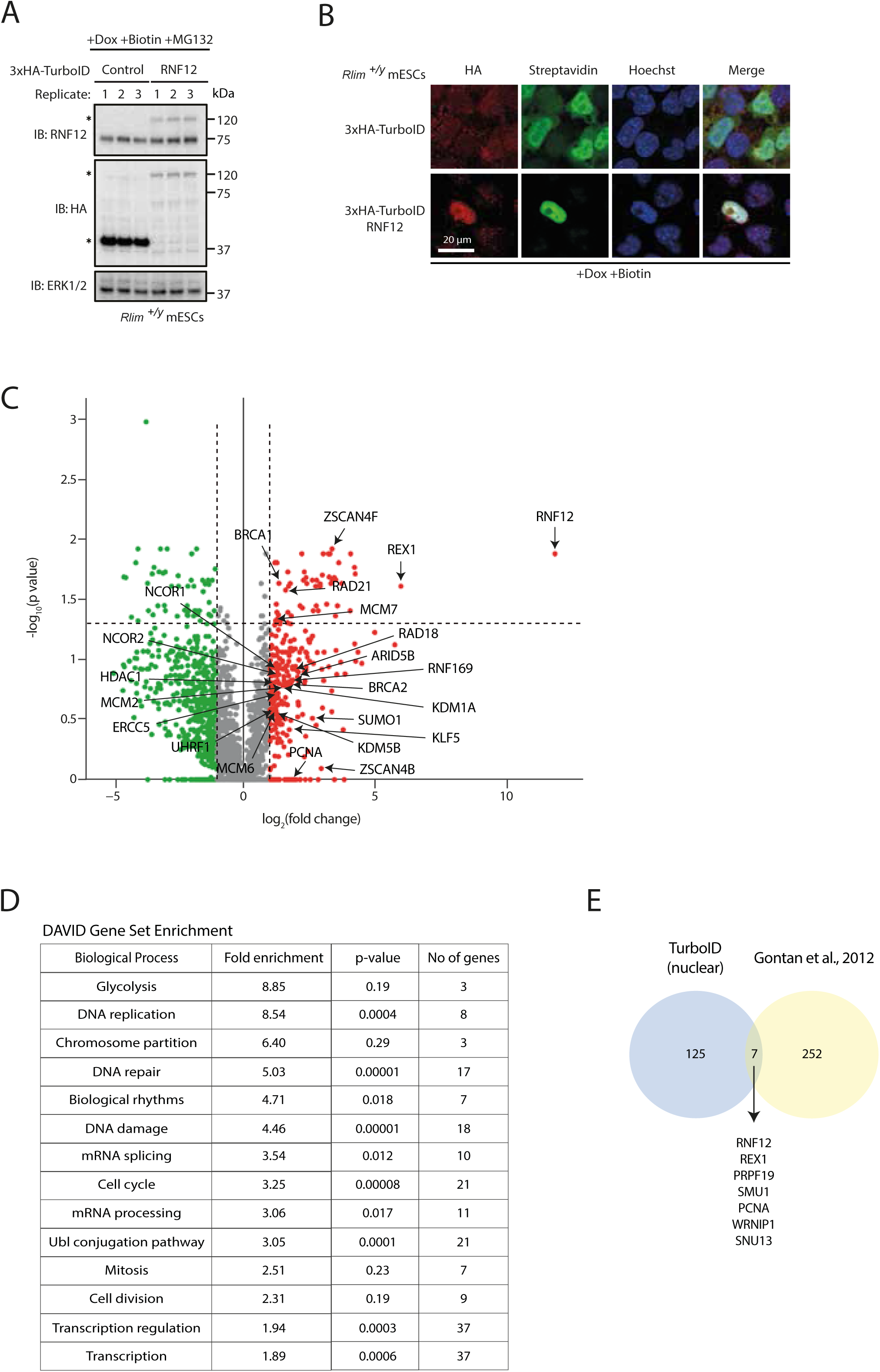
RNF12 TurboID proximity labelling identifies chromatin-associated proteins. (A) Immunoblot analysis of MG132, Doxycycline and Biotin treated *Rlim^+/y^*mESCs stably overexpressing HA-TurboID RNF12 and HA-TurboID control in triplicate. HA-TurboID RNF12 and HA-TurboID control are indicated, ERK1/2 is shown as a loading control (B) Immunofluorescence analysis of Doxycycline and Biotin treated *Rlim^+/y^* mESCs stably overexpressing HA-TurboID RNF12 and HA-TurboID control. Streptavidin staining is shown to confirm protein biotinylation, Hoechst is shown as a nuclear stain. (C) Volcano plot showing relative change in protein abundance of biotinylated proteins comparing MG132 treated mESCs stably overexpressing HA-TurboID RNF12 and HA-TurboID control. Red data points indicate proteins that showed a >2-fold increase in intensity in HA-TurboID RNF12 expressing mESCs. (D) DAVID analysis of enriched genes, grouped by biological process, of those found to have a >2-fold increase in intensity in HA-TurboID RNF12 overexpressing cells and whose translated proteins are annotated with nuclear localisation and/or function. (E) Venn diagram displaying the number of proteins identified by TurboID as having a >2-fold change in expression in *Rlim^+/y^*mESCs overexpressing WT RNF12 relative to control, compared to the number of proteins identified in RNF12 affinity-purification mass spectrometry (Gontan et al., 2012). Proteins common to both datasets are labelled.

Next, we interrogated RNF12 priority proximity labelled proteins for further information about RNF12 regulation and/or function. As RNF12 is localised to the nucleus (Fig. 1B) (Bustos et al., 2020; Jiao et al., 2013), proteins with annotated nuclear localisation and/or function were prioritised from the >2-fold enriched cohort (132 proteins; Table S2). We then performed DAVID gene set enrichment analysis (Huang, Sherman, & Lempicki, 2009; Sherman et al., 2022), which indicates that RNF12 proximity labelled proteins are significantly enriched for chromatin-dependent functions, such as DNA damage response, regulation of gene expression and DNA replication (Fig. 1D). We also compared our findings using TurboID with previously published findings from RNF12 affinity-purification mass-spectrometry (Gontan et al., 2012) (Table S3). Interestingly, in addition to known substrate REX1, only a further five common proteins were identified (Fig. 1E), suggesting that the TurboID approach is able to reveal previously undiscovered RNF12 proximal proteins. Of the common hits, several are chromatin associated including PCNA, SMU1 and WRNIP1. In summary, our data suggest that the chromatin environment may form a key component of RNF12 regulation and function, consistent with our previous data indicating that RNF12 is recruited to chromatin in mESCs (Segarra-Fas et al., 2022).

### RNF12 engages chromatin

In light of these findings, we explored the biological function of RNF12 chromatin recruitment. Using biochemical fractionation, we determined that RNF12 is present in both soluble cytoplasm and/or nucleoplasm, and on chromatin alongside REX1 (Fig. 2A). Effective separation of chromatin from other soluble nuclear/cytoplasmic material was confirmed by immunoblotting for βIII tubulin (TUBβ3), a component of microtubules, and Histone H3 pSer10 (pH3), a core component of chromatin (Fig. 2A). Quantification of the relative amounts of RNF12 and REX1 found on chromatin compared to other cellular locations indicates that a significant proportion of RNF12 (25.7% ± 11.4), and REX1 (52.4% ± 18.8) is recruited to chromatin (Fig. 2B,C). RNF12 (Bustos et al., 2020; Jiao et al., 2013) and REX1 (Gontan et al., 2012) are both localised to the nucleus, suggesting that the remainder is largely present in the nucleoplasm and/or other nuclear structures. These data therefore indicate that RNF12 is recruited to chromatin along with key substrate REX1, although the majority of RNF12 and a significant proportion of REX1 is found in other nuclear compartments.

**Fig. 2.**
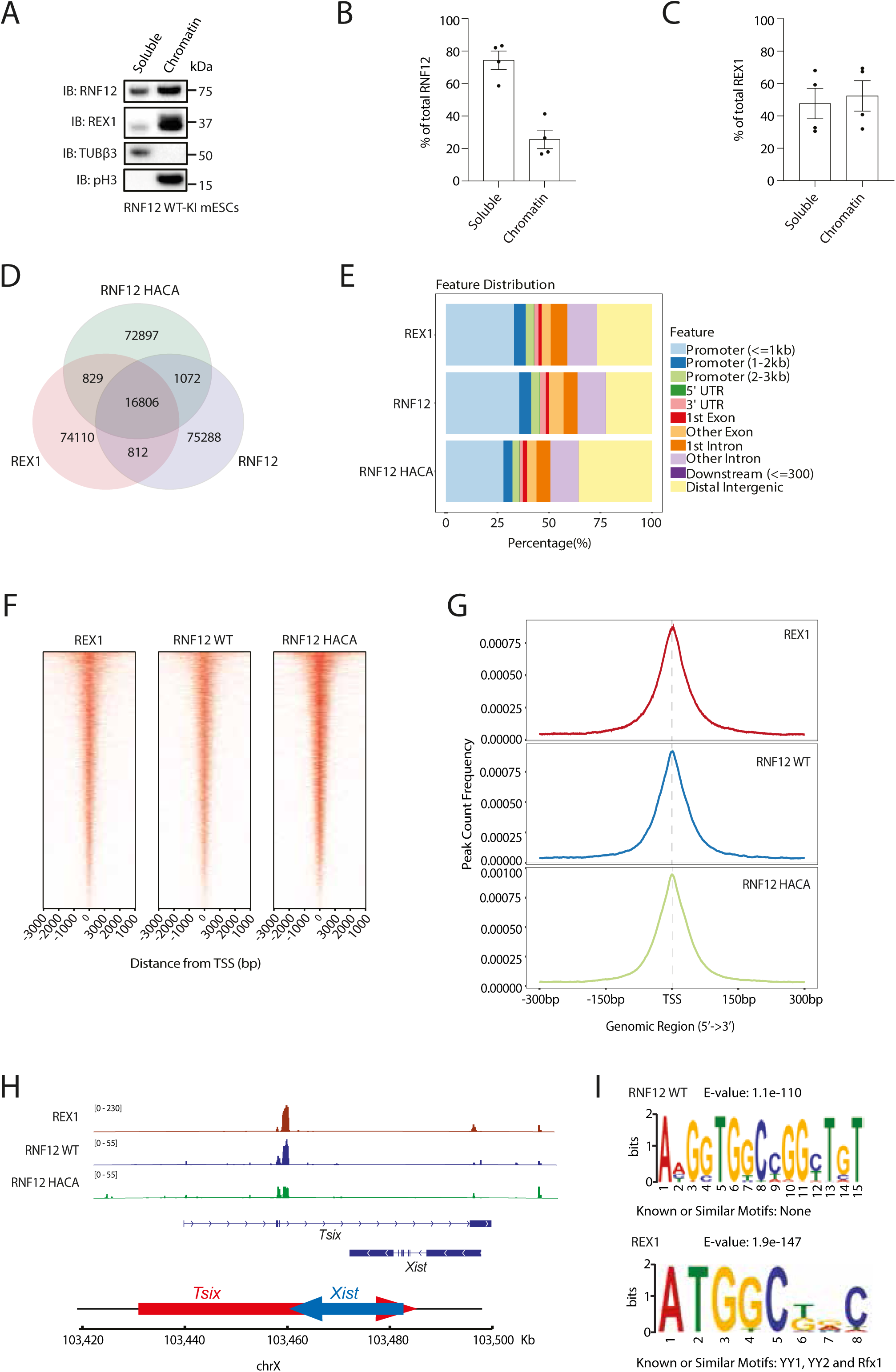
RNF12 colocalises with REX1 transcription factor substrate at specific genomic regions. (A) RNF12 wild-type knock-in (WT-KI) mESCs were subjected to a chromatin fractionation protocol and RNF12, REX1, Tubulin β-3 and phospho-Ser10 Histone H3 (pH3) levels analyzed by immunoblotting. Tubulin β-3 (TUBβ3) is used as a marker of the soluble fraction, and pH3 as a marker of the chromatin fraction. Data are representative of n=4 independent experiments. (B) Graph representing the quantification of the RNF12 signal observed in A. Data is represented as the proportion of RNF12 in soluble and chromatin fractions. As the protein content of the soluble fraction is much higher than that of the chromatin fraction, the proportion of protein in soluble versus chromatin fraction was calculated in all subsequent quantifications. Data represented as mean ± SEM (n=4). (C) Graph representing the quantification of the REX1 signal observed in A. Data represented as mean ± SEM (n=4). (D) Venn diagram showing distribution of overlapping genes with ChIP-seq peaks identified for RNF12, RNF12 catalytically inactive mutant (RNF12^H569A,C572A^) and REX1. (E) Genomic feature distribution of ChIP-seq peaks identified for RNF12, RNF12^H569A,C572A^ and REX1. (F) ChIP-seq peaks identified for RNF12, RNF12^H569A,C572A^ and REX1 clustered according to distance from transcriptional start site (TSS). (G) Peak count frequency relative to distance from transcriptional start sites (TSS) of ChIP-seq peaks identified for RNF12, RNF12^H569A,C572A^ and REX1. (H) Assigned peaks for RNF12, RNF12^H569A,C572A^ and REX1 at the *Xist* gene locus. (I) DNA sequence motif enrichment analysis of ChIP-seq sequences identified for RNF12 and REX1.

### RNF12 and REX1 substrate are co-localised to specific gene regulatory regions

As RNF12 and REX1 are both located on chromatin, we next investigated the genomic regions occupied by these proteins. REX1 genome occupancy in mESCs has been determined previously (Gontan et al., 2012). Therefore, we sought to investigate whether RNF12 and REX1 occupy specific and/or common locations on chromatin by Chromatin Immunoprecipitation (ChIP) followed by DNA sequencing (ChIP-seq). Undifferentiated female WT mESCs treated with the proteasome inhibitor MG132 and expressing FLAG–V5-RNF12 WT and FLAG–V5-RNF12^H569A,C572A^, a catalytically inactive mutant of RNF12 that disrupts the RING domain were employed to perform parallel ChIP-SEQ analyses of RNF12 WT, RNF12^H569A,C572A^ and REX1. Peak assignment was performed to identify associated genome sequence occupancy. We first addressed whether RNF12 genome occupancy overlaps with that of REX1 in mESCs. Although RNF12, RNF12^H569A,C572A^ and REX1 are each recruited to unique chromatin regions, this analysis reveals that RNF12, RNF12^H569A,C572A^ and REX1 are also recruited to shared genome sequences (Fig. 2D). Feature distribution analysis of RNF12, RNF12^H569A,C572A^ and REX1 bound genomic regions indicates an enrichment of promoter proximal regions (Fig. 2E). Furthermore, analysis of the position of RNF12, RNF12^H569A,C572A^ and REX1 chromatin binding sites (Fig. 2F) indicates strong enrichment of RNF12 and REX1 at transcriptional start sites (Fig. 2G). As REX1 was previously shown to be enriched at genomic regions close to transcriptional start sites (Gontan et al., 2012), this is consistent with the overlap observed between RNF12 and REX1 chromatin binding sites. As an example, the region encompassing the long non-coding RNA *Xist* and its antisense transcript *Tsix*, which are located in the X inactivation centre and play crucial roles in the regulation of X chromosome inactivation (Barakat et al., 2011; Gontan et al., 2012; Gontan et al., 2018; Jonkers et al., 2009; Feng Wang et al., 2017), exhibits RNF12 and REX1 peaks of genome occupancy (Fig. 2H). Taken together, our data suggest that RNF12 and REX1 occupy common sites within the genome, in addition to genomic regions that are unique to RNF12 or REX1.

These findings suggest that regional specificity is somehow conferred upon RNF12 genome recruitment. We thus sought to determine the sequence motifs occupied by RNF12 and REX1 within the genome. Analysis of REX1 genomic binding sites suggests enrichment of a DNA sequence motif previous associated with the REX1/YY1/YY2 family of transcriptional regulators (Fig. 2I) (J. D. Kim, Faulk, & Kim, 2007). Interestingly, analysis of RNF12 recruitment sites, which exhibit some overlap with REX1 recruitment sites, reveals a distinct sequence recruitment motif (Fig. 2I). These findings indicate that although RNF12 and REX1 are recruited to shared genomic sites, this may occur via distinct sequence motifs.

### RNF12 substrate REX1 is efficiently ubiquitylated specifically on chromatin

Our demonstration that RNF12 and REX1 are co-located on specific gene regulatory regions prompts the hypothesis that chromatin recruitment is a key event to enable RNF12 ubiquitylation of REX1 at specific genomic locations to regulate gene expression. Therefore, we measured REX1 ubiquitylation on chromatin and in other cellular compartments by stabilising ubiquitylated REX1 using the proteasomal inhibitor MG132 and performing chromatin fractionation. This analysis suggests that endogenous REX1 is heavily ubiquitylated in the chromatin fraction compared to other cellular compartments (Fig. 3A). As expected, REX1 ubiquitylation is reduced in RNF12-deficient mESCs (*Rlim*^-/y^), although residual REX1 ubiquitylation is observed, particularly shorter chains. Quantification indicates that the majority of REX1 ubiquitylation occurs on chromatin (Fig. 3B), and this is reduced is RNF12-deficient mESCs (Fig. 3C). RNF12-dependent REX1 ubiquitylation on chromatin was also increased in RNF12-deficient mESCs reconstituted with RNF12 WT, but not with a catalytic-deficient mutant of RNF12 (RNF12 W576Y) (Fig 3D,E), indicating that enriched REX1 ubiquitylation on chromatin requires RNF12 catalytic activity.

**Fig. 3.**
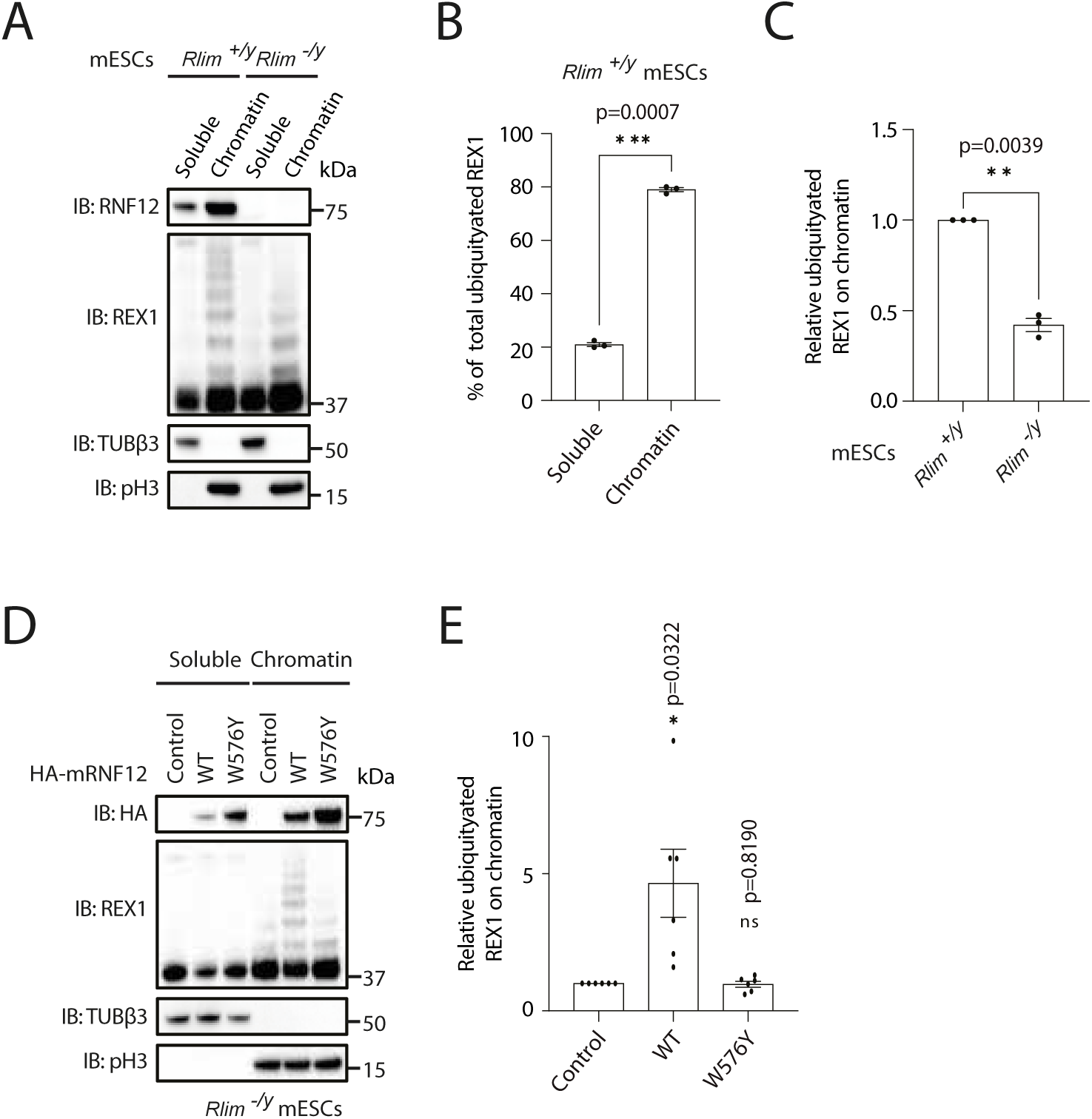
RNF12 substrate REX1 is efficiently ubiquitylated on chromatin. (A) Immunoblotting for REX1 ubiquitylation levels in control (*Rlim^+/y^*) and RNF12 knock-out (*Rlim^-/y^*) mESCs subjected to chromatin fractionation. Cells were treated with MG132 inhibitor for 1 h to allow visualization of ubiquitylated REX1. Data are representative of n=3 independent experiments. (B) REX1 ubiquitylation signal in soluble and chromatin fractions in *Rlim^+/y^* mESCs observed in A was quantified and normalized to total REX1 levels. Data represented as mean ± SEM (n=3). Statistical significance was determined by paired *t* test; two-sided, confidence level 95%. (C) REX1 ubiquitylation signal on chromatin in *Rlim^+/y^*and *Rlim^-/y^* mESCs was quantified and normalized to total REX1 levels. Data represented as mean ± SEM (n=3). Statistical significance was determined by paired *t* test; two-sided, confidence level 95%. (D) Immunoblotting for REX1 ubiquitylation levels in *Rlim^-/y^* mESCs expressing either empty vector (control), HA-mRNF12 WT and HA-mRNF12 W576Y and subjected to a chromatin fractionation protocol. *Rlim^-/y^* mESCs were treated with MG132 for 1 h to allow visualization of ubiquitylated REX1. Data are representative of n=6 independent experiments. (E) REX1 ubiquitylation signal on chromatin was quantified and normalized to total REX1 levels. Data represented as mean ± SEM (n=6). Statistical significance was determined by paired *t* test; two-sided, confidence level 95%.

### RNF12 is recruited to chromatin via the Basic Region (BR)

As the majority of RNF12-dependent REX1 ubiquitylation take places on chromatin, we next sought to address the mechanism by which RNF12 engages chromatin. To this end, we performed deletion mutagenesis to identify RNF12 sequences that are required for chromatin recruitment (Fig. 4A). As expected, deletion of the RNF12 Nuclear Localization Signal (NLS) reduced chromatin recruitment (Fig. 4B,C), presumably via effects on RNF12 nuclear localisation. In contrast, deletion of the RNF12 Nuclear Export Signal (NES) or the catalytic RING domain has no discernible impact on chromatin recruitment (Fig. 4B,C). However, Basic Region (BR) deletion prevents recruitment to chromatin, suggesting that the basic region is a major determinant of RNF12 chromatin recruitment. Interestingly, deletion of the N-terminal sequences drives increased RNF12 chromatin recruitment (Fig. 4B,C), suggesting that the RNF12 N-terminus functions to inhibit RNF12 chromatin engagement.

**Fig. 4.**
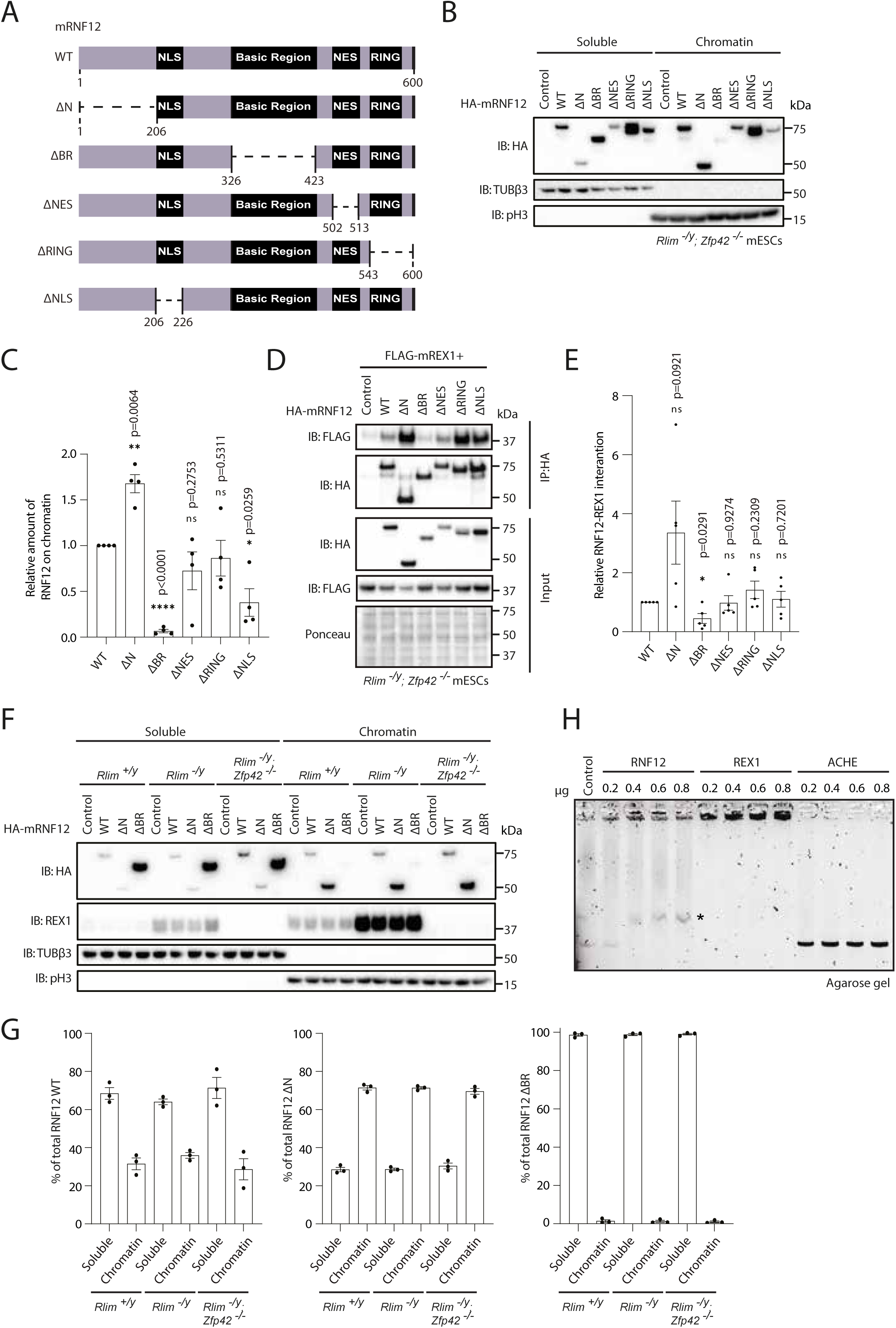
RNF12 chromatin recruitment is largely REX1 independent. (A) Schematic of the structure of mouse RNF12 WT and deletion mutants. Indicated are the amino acid boundaries of each deletion. (B) Immunoblotting for HA-tagged mouse RNF12 WT (1-600), RNF12 Δ1-206 (ΔN), RNF12 Δ326-423 (ΔBR), RNF12 Δ502-513 (ΔNES), RNF12 Δ543-600 (ΔRING) and RNF12 Δ206-226 (ΔNLS) in soluble and chromatin fractions of transfected RNF12 and REX1 double knock-out (*Rlim^-/y^; Zfp42^-/-^*) mESCs. Data are representative of n=4 independent experiments. (C) Graph representing the quantification of the HA-RNF12 signal observed in the chromatin fraction in B. The percentage of each deletion mutant in the chromatin fraction was normalized to that observed for RNF12 WT. Data represented as mean ± SEM (n=4). Statistical significance was determined by paired *t* test; two-sided, confidence level 95%. (D) Immunoblotting for HA and FLAG in HA immunoprecipitates from RNF12 and REX1 double knock-out (*Rlim^-/y^; Zfp42^-/-^*) mESCs expressing FLAG-REX1 with either empty vector (control) or the indicated HA-mRNF12 deletions mutants. RNF12 and REX1 double knock-out (*Rlim^-/y^; Zfp42^-/-^*) were treated with MG132 for 2 h and Ponceau S staining is shown as a loading control. Data are representative of n=5 independent experiments. (E) Graph representing the quantification of experiments from D. Data represented as mean ± SEM (n=5). Statistical significance was determined by paired *t* test; two-sided, confidence level 95%. (F) HA-tagged mouse RNF12 WT, RNF12 ΔN and RNF12 ΔBR levels in soluble and chromatin fractions were analysed by immunoblotting in control (*Rlim^+/y^*), RNF12 knock-out (*Rlim^-/y^*) and RNF12 and REX1 double knock-out (*Rlim^-/y^; Zfp42^-/-^*) mESCs. REX1, Tubulin β-3 and phospho-S10 H3 levels are also shown. Data are representative of n=3 independent experiments. (G) Graph representing the quantification of the HA-tagged mouse RNF12 signal observed in F. Data represented as mean ± SEM (n=3). (H) Electrophoretic mobility shift analysis (EMSA) of linearized pCAGGS plasmid DNA (0.5 µg) incubated with increasing concentrations (0.2-0.8 µg) of RNF12, REX1 and ACHE recombinant proteins. An 0.8% agarose gel was used. Data are representative of n=3 independent experiments.

### RNF12 chromatin recruitment mechanism is REX1 independent

Previous work has shown that the RNF12 BR is required for REX1 interaction (Gontan et al., 2012), suggesting that substrate engagement may be a key mechanism for chromatin recruitment. We confirmed that the RNF12 BR is required for interaction with REX1 substrate. In immunoprecipitation assays, RNF12 interacts with REX1 and this is reduced by deletion of the RNF12 BR (Fig. 4D,E), confirming the role of the RNF12 BR in REX1 substrate interaction. Interestingly, RNF12 N-terminal deletion leads to increased REX1 binding (Fig. 4D,E), suggesting that the RNF12 N-terminus not only inhibits chromatin recruitment, but also REX1 substrate interaction.

Considering our finding that the RNF12 Basic and N-terminal regions mediate both chromatin recruitment and REX1 interaction, we tested whether REX1 engagement is responsible for RNF12 chromatin recruitment. To this end, we took advantage of an allelic series of wild-type (WT), RNF12-deficient (*Rlim*^-/y^) and RNF12/REX1-deficient (*Rlim*^-/y^; *Zfp42*^-/-^) mESC lines reconstituted with HA-RNF12 WT. HA-RNF12 is efficiently recruited to chromatin in either an RNF12-deficient or RNF12/REX1-deficient background (Fig. 4F,G), suggesting that interaction with REX1 is not a major mechanism for RNF12 chromatin recruitment. Consistent with this notion, recruitment of RNF12 BR and N-terminal deletion mutants to chromatin is not altered by REX1 deletion (Fig. 4F,G).

Our data indicating that REX1 is largely dispensable for RNF12 chromatin recruitment suggest that the major mechanism of RNF12 chromatin engagement is independent of REX1 interaction. As RNF12 chromatin recruitment is mediated by the BR, we asked whether this positively charged region might mediate direct electrostatic interactions with negatively charged DNA. To test this, we incubated recombinant RNF12 with circular plasmid DNA (pCAGGS) and performed electrophoretic mobility shift analysis (EMSA). In the absence of protein or in the presence of negative control protein ACHE, pCAGGS plasmid is resolved at the expected molecular weight by agarose gel electrophoresis (Fig. 4H). However, addition of the REX1 transcription factor, which directly binds DNA, reduces the electrophoretic mobility of plasmid DNA upon EMSA (Fig. 4H). Similarly, RNF12 reduces the electrophoretic mobility of plasmid DNA upon EMSA (Fig. 4H, asterisk), suggesting that RNF12 also has the capacity to directly interact with DNA. Taken together, our results indicate that the RNF12 BR mediates recruitment to chromatin in a manner that is independent of REX1 interaction, potentially by directly interacting with DNA.

### RNF12 chromatin recruitment via the Basic Region is required for substrate processing

We next sought to determine the specific sequences within the RNF12 BR that are required for chromatin recruitment. To this end, we generated three smaller BR deletions (ΔBR1 lacking amino acids 326-348, ΔBR2 lacking amino acids 348-381 and ΔBR3 lacking amino acids 381-423) (Fig. 5A) and addressed the impact of these sequences on chromatin recruitment. As shown previously, deletion of the RNF12 BR abolishes chromatin recruitment (Fig. 5B,C). Similarly, deletion of BR1 and BR2 disrupts chromatin recruitment, although to a lesser extent (Fig. 5B,C). In contrast, deletion of BR3 increases RNF12 chromatin recruitment (Fig. 5B,C). This region has fewer basic residues than BR1 and BR2, suggesting that BR3 is not only dispensable for chromatin engagement, but may encode an element that autoinhibits engagement of chromatin by RNF12.

**Fig. 5.**
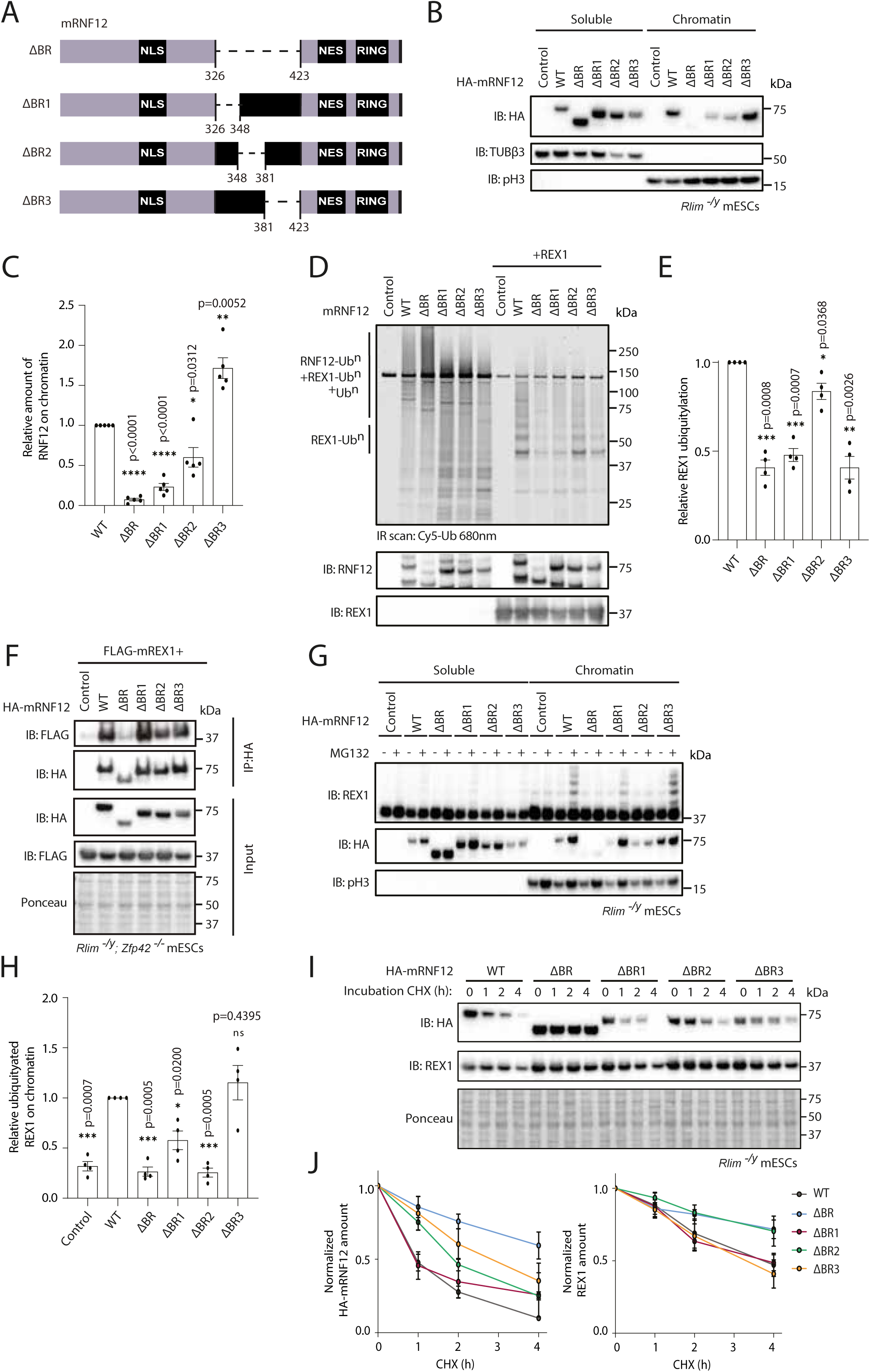
RNF12 chromatin recruitment and substrate ubiquitylation is mediated by the Basic Region (BR) (A) Schematic representation of mouse RNF12 BR, BR1, BR2 and BR3 deletions. (B) Immunoblotting for HA-tagged mRNF12 on soluble and chromatin fraction of transfected RNF12 knock-out (*Rlim^-/y^*) mESCs with HA-tagged versions of RNF12 WT and RNF12 BR deletions. Data are representative of n=5 independent experiments. (C) Graph representing the quantification of the HA-mRNF12 signal observed in the chromatin fraction in B. The percentage of each deletion mutant in the chromatin fraction was normalized to that observed for RNF12 WT. Data represented as mean ± SEM (n=5). Statistical significance was determined by paired *t* test; two-sided, confidence level 95%. (D) *In vitro* REX1 substrate ubiquitylation assay of RNF12 WT and RNF12 BR deletions. Top: Fluorescently labelled ubiquitylated proteins were detected by scan at 680nm (Cy5-Ub) after 30 mins of reaction, and specific ubiquitylated REX1 (REX1-Ub^n^) and RNF12 (RNF12-Ub^n^) signals are indicated. Bottom: Immunoblot analysis of RNF12 (using anti-RNF12 mouse monoclonal antibody) and REX1 protein levels. Data are representative of n=4 independent experiments. (E) REX1 ubiquitylation signal was quantified and normalized to total REX1 amounts. Only the first three ubiquitylated bands (REX1-Ub^3^) were quantified. Data represented as mean ± SEM (n=4). Statistical significance was determined by paired *t* test; two-sided, confidence level 95%. (F) Immunoblotting for HA and FLAG in HA immunoprecipitates from RNF12 and REX1 double knock-out (*Rlim^-/y^; Zfp42^-/-^*) mESCs expressing FLAG-REX1 with either empty vector (control) or the indicated HA-RNF12 deletions mutants. RNF12 and REX1 double knock-out (*Rlim^-/y^; Zfp42^-/-^*) mESCs were treated with MG132 inhibitor for 2 h and Ponceau S staining is shown as a loading control. Data are representative of n=4 independent experiments. (G) RNF12 knock-out (*Rlim^-/y^*) mESCs expressing HA-tagged mouse RNF12 WT or BR deletions were treated with either DMSO (vehicle control) or MG132 for 1 h. REX1 ubiquitylation levels are shown. Data are representative of n=4 independent experiments. (H) REX1 ubiquitylation signal on the chromatin fraction in the presence of MG132 in G was quantified and normalized to the signal observed after transfection of HA-mRNF12 WT in RNF12 knock-out (*Rlim^-/y^*) mESCs. Data represented as mean ± SEM (n=4). Statistical significance was determined by paired *t* test; two-sided, confidence level 95%. (I) HA-mouse RNF12 and REX1 half-life was analyzed by immunoblotting in RNF12 knock-out (*Rlim^-/y^*) mESCs expressing HA-tagged mouse RNF12 WT, RNF12 ΔBR, RNF12 ΔBR1, RNF12 ΔBR2 and RNF12 ΔBR3. Cells were treated with 350 µM cycloheximide (CHX) for the indicated times. Ponceau S staining is shown as a loading control. Data are representative of n=4 independent experiments. (J) Representation of the HA-tagged RNF12 and REX1 proteins levels quantified in I. Protein levels were normalized to the Ponceau S loading control. Data represented as mean ± SEM (n=4). Statistical significance of each deletion mutant compared to HA-mRNF12 WT was determined at 2 h for HA-tagged mouse RNF12 and at 4 h for REX1 by paired *t* test; two-sided, confidence level 95%. HA-tagged mRNF12: RNF12 ΔBR (**) p=0.0026, RNF12 ΔBR1 (ns) p=0.6554, RNF12 ΔBR2 (*) p=0.0393 and RNF12 ΔBR3 (*) p=0.0186. REX1: RNF12 ΔBR (*) p=0.0322, RNF12 ΔBR1 (ns) p=0.7590, RNF12 ΔBR2 (**) p=0.0050 and RNF12 ΔBR3 (ns) p=0.2781.

As RNF12 BR deletions can, in principle, impact on catalytic activity (Bustos et al., 2018), REX1 substrate recruitment (Gontan et al., 2012) (Fig. 4D,E) and/or chromatin recruitment (Fig. 4B,C), we set out to distinguish these possibilities. To this end, we measured catalytic activity of RNF12 BR deletion mutants in the presence of recombinant ubiquitin, UBE2D1 (E2), UBE1 (E1) and REX1 substrate *in vitro*. As shown previously, RNF12 WT catalyses REX1 substrate ubiquitylation (Bustos et al., 2018) (Fig. 5D,E). Interestingly, deletion of BR, BR1 or BR3 significantly decreases REX1 ubiquitylation (Fig. 5D,E), although RNF12 catalytic activity towards REX1 is largely unaffected by BR2 deletion (Fig. 5D,E). As expected, engagement of REX1 by RNF12 is largely unaffected by BR2 deletion, when compared to BR deletion (Fig. 5F). These data indicate that whilst the RNF12 BR performs functions that are required for chromatin recruitment, substrate engagement and catalysis, specific deletion of the RNF12 BR2 region separates these functions by impacting primarily on chromatin recruitment, without significantly impacting on catalytic activity and substrate engagement.

We then explored the effect of RNF12 BR deletions on REX1 substrate ubiquitylation in mESCs. Using MG132 treatment in combination with chromatin fractionation as before, we were able to sensitively measure RNF12-dependent REX1 ubiquitylation (Fig. 5G,H). Consistent with the impact on chromatin recruitment, the RNF12 BR is required for efficient REX1 ubiquitylation (Fig. 5G,H). However, RNF12 BR1 and BR3 deletions drive efficient REX1 ubiquitylation (Fig. 5G,H), despite differing relative impacts on chromatin recruitment (Fig. 5B,C). Although REX1 ubiquitylation is observed with RNF12 BR1 deletion, this appears to be a consequence of increased chromatin recruitment in the presence of MG132, when compared to that observed for RNF12 BR2 and BR3 deletions (Fig. 5G). In contrast, RNF12 BR2 deletion is impaired for both REX1 ubiquitylation (Fig. 5G,H) and chromatin recruitment (Fig. 5B,C), suggesting that the BR2 region plays a critical role in RNF12 chromatin recruitment, which in turn impacts on substrate ubiquitylation. This notion is supported by REX1 stability, which is more profoundly affected by RNF12 BR and BR2 deletion, when compared to RNF12 BR1 and BR3 deletion (Fig. 5I,J). However, in contrast to RNF12 BR deletion, RNF12 BR2 deletion mutant undergoes efficient degradation mediated by autoubiquitylation (Fig. 5I,J), consistent with our observation that RNF12 BR2 deletion does not have a major impact on catalytic activity *per se* (Fig. 5D). Therefore, these data support the conclusion that the RNF12 BR is critical for REX1 substrate ubiquitylation by enabling RNF12 recruitment to chromatin.

### RNF12 N-terminal region negatively regulates chromatin recruitment and substrate ubiquitylation

We have demonstrated that the RNF12 BR is required for chromatin recruitment and substrate ubiquitylation/binding. However, we observed an opposing effect of the RNF12 N-terminal region, deletion of which leads to increased RNF12 chromatin association, suggesting that the RNF12 N-terminus somehow acts to suppress chromatin recruitment. This prompted us to address the functional importance of the RNF12 N-terminal region in substrate ubiquitylation.

First, we sought to define the specific sequences within the RNF12 N-terminal region that are required for chromatin recruitment. To this end, we generated three smaller deletions of the RNF12 N-terminus (ΔN1 lacking amino acids 1-68, ΔN2 lacking amino acids 68-135 and ΔN3 lacking amino acids 135-206) to determine the impact of these N-terminal sequences on chromatin recruitment (Fig. 6A). As shown previously, deletion of the RNF12 N-terminus augments chromatin recruitment (Fig. 4B,C). However, deletion of N1, N2 and N3 have no significant impact on RNF12 chromatin recruitment (Fig. 6B,C), and behave similarly to RNF12 WT in these assays. These data suggest that the entire RNF12 N-terminal region (amino acids 1-206) is required to negatively regulate chromatin recruitment.

**Fig. 6.**
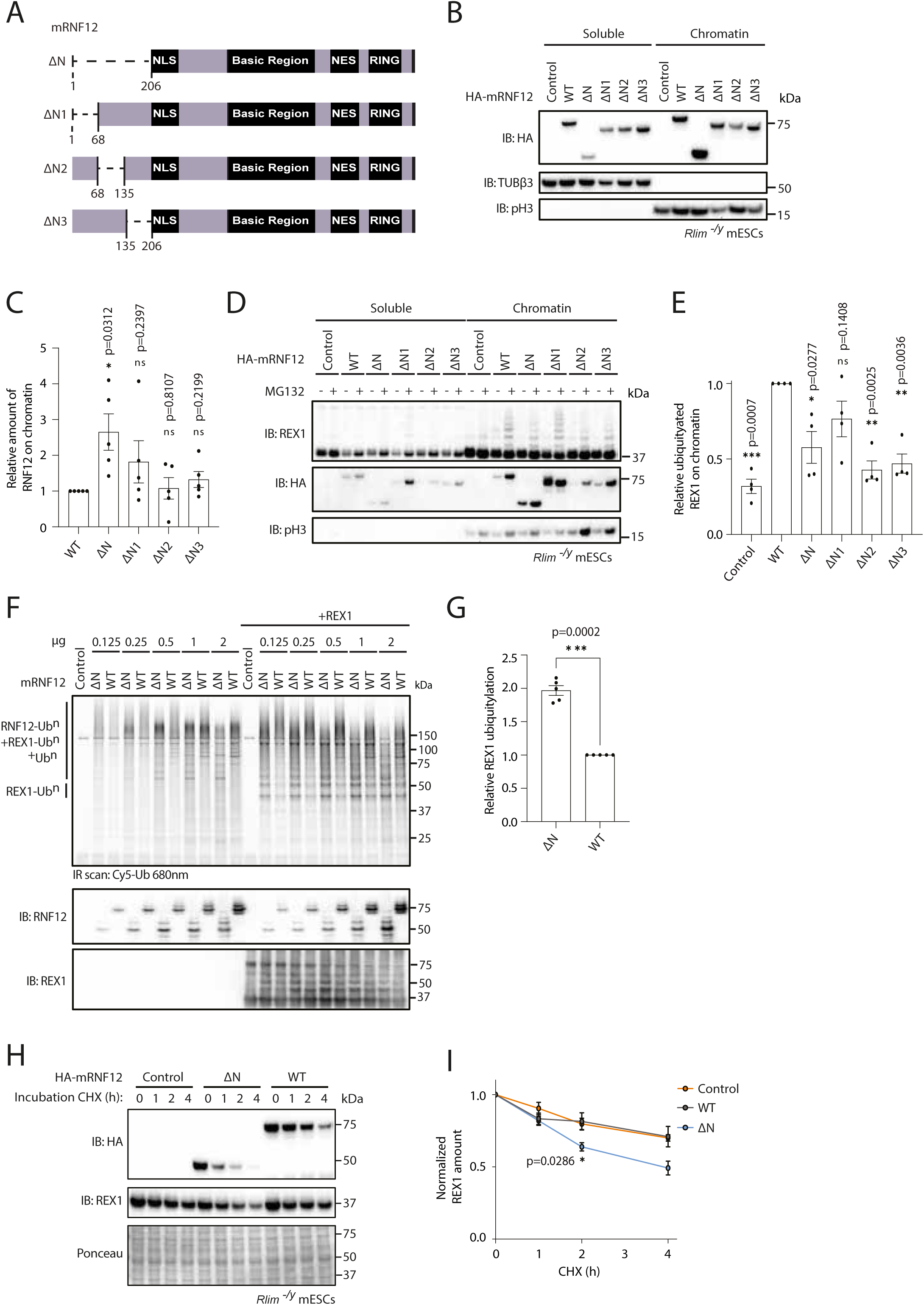
RNF12/RLIM N-terminal region inhibits chromatin recruitment and substrate ubiquitylation. (A) Schematic representation of mouse RNF12 N-terminal deletion (ΔN), and RNF12 N1, N2 and N3 deletions. (B) Immunoblotting for HA-tagged mouse RNF12 in soluble and chromatin fractions of RNF12 knock-out (*Rlim^-/y^*) mESCs expressing HA-tagged mouse RNF12 WT and N-terminal deletions. Data are representative of n=5 independent experiments. (C) Graph representing the quantification of the HA-RNF12 signal observed in the chromatin fraction. The percentage of each deletion mutant in the chromatin fraction was normalised to that observed for mRNF12 WT. Data represented as mean ± SEM (n=5). Statistical significance was determined by paired *t* test; two-sided, confidence level 95%. (D) RNF12 knock-out (*Rlim^-/y^*) mESCs expressing HA-tagged mouse RNF12 WT and N-terminal deletions were treated with MG132 for 1 h. REX1 ubiquitylation levels are shown. Data are representative of n=4 independent experiments. (E) REX1 ubiquitylation signal in the chromatin fraction in the presence of MG132 was quantified and normalized to the signal observed after transfection of HA-RNF12 WT in *Rlim^-/y^*mESCs. Data represented as mean ± SEM (n=4). Statistical significance was determined by paired *t* test; two-sided, confidence level 95%. (F) *In vitro* REX1 substrate ubiquitylation assay of WT mRNF12 and mRNF12 N-terminal deletion at different amounts. Top: Fluorescently labelled ubiquitylated proteins were detected by scan at 680nm (Cy5-Ub) after 30 mins of reaction and specific ubiquitylated REX1 (REX1-Ub^n^) and RNF12 (RNF12-Ub^n^) signals are indicated. Bottom: Immunoblot analysis of RNF12 (using anti-RNF12 mouse monoclonal antibody) and REX1 protein levels. Data are representative of n=5 independent experiments. (G) REX1 ubiquitylation signal was quantified and normalized to total REX1 amounts. Only the first three ubiquitylated bands (REX1-Ub^3^) were quantified. Data represented as mean ± SEM (n=4). Statistical significance was determined by paired *t* test; two-sided, confidence level 95%. (H) REX1 half-life was analyzed by immunoblotting of RNF12 knock-out (*Rlim^-/y^*) mESCs expressing either empty vector (control), HA-mRNF12 WT and HA-mRNF12 N-terminal deletion. Cells were treated with 350 µM cycloheximide (CHX) for the indicated times. Ponceau staining is shown as a control. Data are representative of n=5 independent experiments. (I) Representation of the REX1 proteins levels quantified in H. Protein levels were normalized to the Ponceau control. Data represented as mean ± SEM (n=5). Statistical significance of REX1 stability in *Rlim^-/y^* mESCs expressing HA-mRNF12 N-terminal deletion compared to those expressing HA-mRNF12 WT was determined at 2 h by paired *t* test; two-sided, confidence level 95%.

We then explored the functional impact of RNF12 N-terminal sequences on REX1 substrate ubiquitylation. Using MG132 treatment in combination with chromatin fractionation, we measured REX1 ubiquitylation as before. These data suggest that the RNF12 N-terminus is required for REX1 ubiquitylation (Fig. 6D,E), in contrast with previous data indicating that the RNF12 N-terminus suppresses REX1 substrate recruitment (Fig. 4D). To resolve these apparently contradictory results, we investigated the direct impact of the RNF12 N-terminal region on E3 ubiquitin ligase activity in the presence of recombinant ubiquitin, UBE2D1 (E2) and UBE1 (E1) and REX1 *in vitro*. As shown previously (Fig. 5D), RNF12 WT specifically ubiquitylates REX1 substrate *in vitro* (Fig. 6F,G). Deletion of the RNF12 N-terminal region significantly increases REX1 ubiquitylation (Fig. 6F,G), suggesting that the RNF12 N-terminus suppresses chromatin recruitment, substrate engagement and inhibits catalysis. Consistent with these impacts, RNF12 N-terminal deletion augments REX1 degradation in cells, under conditions where RNF12 WT levels are limiting for REX1 processing (Fig. 6H,I). Taken together, these data indicate an apparently autoinhibitory function of the RNF12 N-terminal region in suppressing chromatin recruitment, REX1 substrate recruitment and ubiquitylation.

### RNF12 chromatin recruitment is required for patterning gene expression

Finally, we sought to determine the functional importance of RNF12 chromatin recruitment for patterning of developmental gene expression. One of the key functions of RNF12 during development is induction of imprinted X-chromosome inactivation (Barakat et al., 2011; Gontan et al., 2018; Shin et al., 2010; Feng Wang et al., 2016), which occurs by relieving REX1-mediated transcription repression of the long-non-coding RNA *Xist* via REX1 ubiquitylation and degradation (Barakat et al., 2011; Gontan et al., 2012). Therefore, we used an assay for ectopic *Xist* induction by RNF12 expression in male mESCs (Barakat et al., 2011; Gontan et al., 2012; Gontan et al., 2018; Jonkers et al., 2009; Shin et al., 2010), which serves as a sensitive readout of RNF12-dependent transcriptional responses mediated by REX1 degradation. As expected, *Xist* expression is low in male mESCs, but expression of HA-RNF12 WT drives *Xist* induction (Fig. 7A,B). In contrast, expression of catalytic deficient RNF12 W576Y fails to drive *Xist* induction (Fig. 7A,B), indicating that *Xist* induction is dependent upon RNF12 E3 ubiquitin ligase activity. Similarly, expression of RNF12 deletions lacking either the entire BR or the BR2 region that impacts largely on chromatin recruitment but not on substrate binding or catalysis, fails to drive *Xist* induction (Fig. 7A,B). These data therefore provide evidence that chromatin recruitment of RNF12 by the BR plays a key role in patterning of developmental gene expression, as measured by *Xist* induction.

**Fig. 7.**
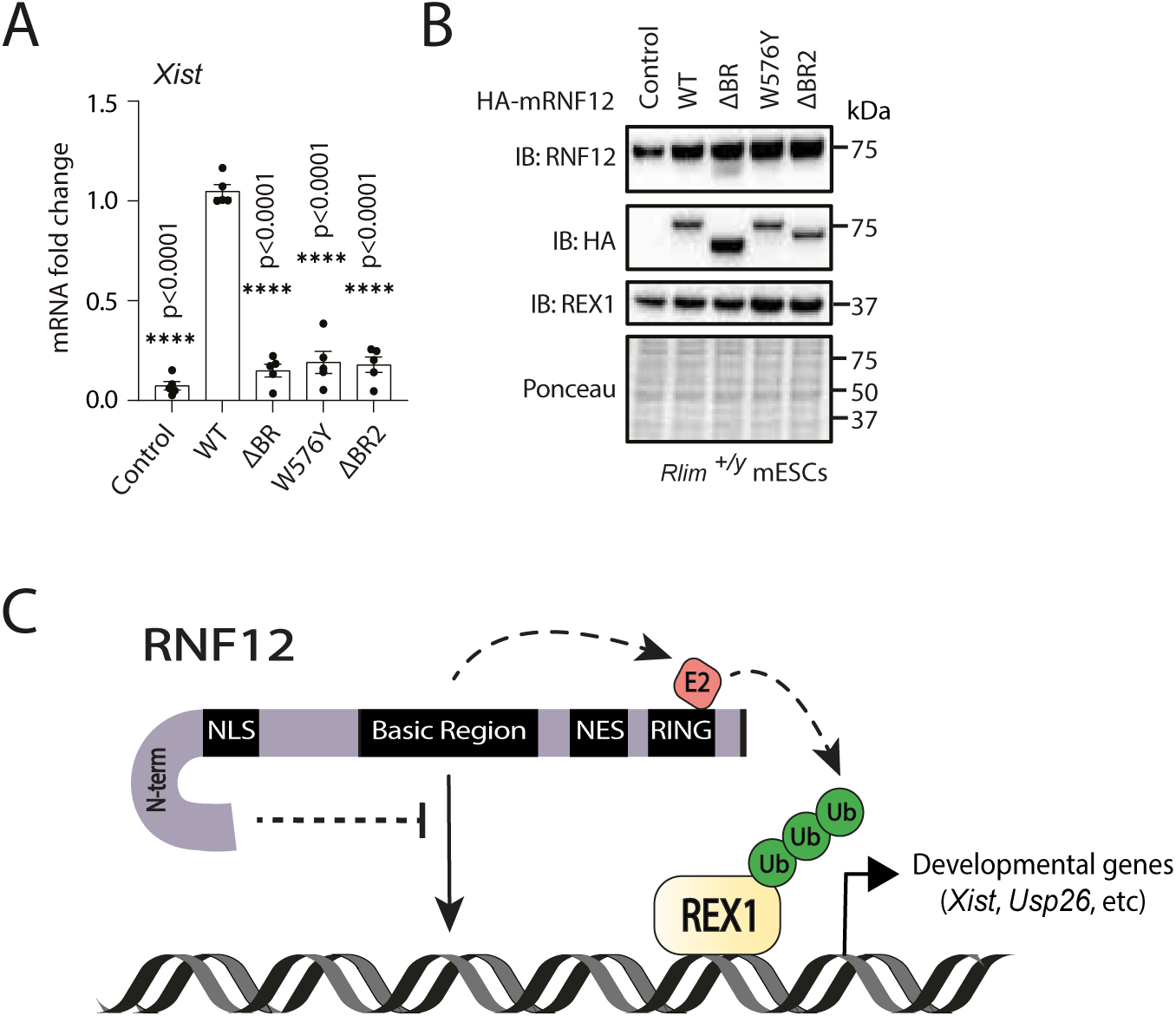
RNF12 chromatin recruitment is required for transcriptional patterning. (A) Quantification of *Xist* mRNA in *Rlim^+/y^* mESCs expressing HA-tagged mouse RNF12 WT, RNF12 BR deletion, RNF12 W576Y and RNF12 BR2 deletion by qRT-PCR analysis. *Xist* lncRNA expression was normalised to *Gapdh* mRNA expression. Data represented as mean ± SEM (n=5). Statistical significance was determined by paired *t* test; two-sided, confidence level 95%. (B) RNF12, HA and REX1 expression was analysed by immunoblotting in *Rlim^+/y^* mESCs expressing HA-tagged versions of mouse RNF12 WT, RNF12 BR deletion, RNF12 W576Y and RNF12 BR2 deletion. Ponceau staining is shown as a control. Data are representative of n=5 independent experiments. (C) Model for how chromatin functions as an RNF12 regulatory platform. RNF12 recruitment to chromatin is mediated by RNF12 Basic Region, which is required for efficient REX1 ubiquitylation and regulation of RNF12-dependent genes. In an opposing manner, the RNF12 N-terminal region supresses chromatin recruitment and substrate ubiquitylation, conferring a previously unappreciated autoinhibitory mechanism.

## Discussion

In this paper, we uncover a critical role for chromatin in regulation of substrate ubiquitylation and downstream regulation of gene expression by the RING type E3 ubiquitin ligase RNF12. We show that RNF12 is recruited to chromatin in a spatially restricted manner, including co-recruitment at sites occupied by key substrate, the transcription factor REX1. We show that recruitment to these specific locations facilitates RNF12 mediated ubiquitylation of REX1, thereby patterning expression of RNF12-REX1 dependent genes, including the key long-non-coding RNA *Xist* and its antisense transcript *Tsix*, which collectively coordinate X-chromosome inactivation. Furthermore, we unveil the mechanism underpinning chromatin recruit, whereby a conserved RNF12 basic region (BR) of RNF12 independent of the catalytic RING domain is absolutely required. In addition, we show that the RNF12 BR and another non-RING element at the N-terminal region perform key regulatory functions on chromatin. RNF12 BR is required for chromatin recruitment, and substrate engagement and ubiquitylation, whilst the N-terminal region performs an autoinhibitory function, which prevents chromatin recruitment, substrate engagement and ubiquitylation (Fig. 7C). In combination, this system is required for RNF12 substrate ubiquitylation and regulation of gene expression, providing insight into mechanisms for spatial coordination of substrate ubiquitylation to ensure specific biological outcome.

Yet to be resolved is structural detail of how autoinhibition, chromatin recruitment, substrate engagement and ubiquitylation are coordinated. Although the RNF12 BR is required for chromatin recruitment, REX1 engagement and ubiquitylation, REX1 itself does not mediate the majority of RNF12 chromatin recruitment, suggesting that these mechanisms are separable. However, REX1 may be required to recruit RNF12 to specific genomic locations. Indeed, the RNF12 BR appears to make distinct contacts with chromatin and REX1, which together facilitate REX1 ubiquitylation on chromatin. Furthermore, the mechanism by which the N-terminal region inhibits chromatin recruitment and ubiquitylation is not yet known. However, our data suggests that the N-terminus makes autoinhibitory contacts to inhibit chromatin interaction and substrate engagement/ubiquitylation. Indeed, Alphafold predictions indicate that the RNF12 N-terminal region forms direct contacts with the BR, which may in turn occlude chromatin and/or substrate interaction sites. Whether this negative regulatory system is released by chromatin recruitment or by another signal remains to be determined. Interestingly, there are phosphorylation sites in proximity to the RNF12 N-terminal region that may modulate chromatin engagement and/or substrate ubiquitylation.

In this study, we also reveal that RNF12 is recruited to a specific genomic consensus motif, which potentially confers spatial regulation of REX1 ubiquitylation at specific genomic locations. In that regard, we show that the majority of REX1 ubiquitylation occurs on chromatin, presumably at sites of RNF12 co-recruitment. This prompts the exciting hypothesis that REX1 is specifically ubiquitylated by RNF12 at gene regulatory elements. However, although REX1 appears to play a key role as an accessory factor, it is not required for global recruitment of RNF12 to chromatin, suggesting the existence of other factors that determine sites of chromatin engagement. Future work will explore whether transcriptional components other than REX1 play a role, or whether RNF12 encodes capacity to directly engage chromatin via its specific DNA binding consensus.

Finally, as RNF12 is mutationally disrupted in patients with the X-linked intellectual disability disorder Tonne-Kalscheuer syndrome (TOKAS), an attractive hypothesis states that RNF12 chromatin recruitment and the regulatory systems uncovered herein may be impacted by TOKAS variants. RNF12 TOKAS patient variants are found largely clustered in the catalytic RING domain or the BR. Therefore, a priority will be to determine the impact of RNF12 BR mutations on chromatin recruitment, or whether these largely impact on catalysis, as has been previously shown (Bustos et al., 2018).

## MATERIALS AND METHODS

A full summary of reagents used in this study can be found in Table 1

**Table 1:**
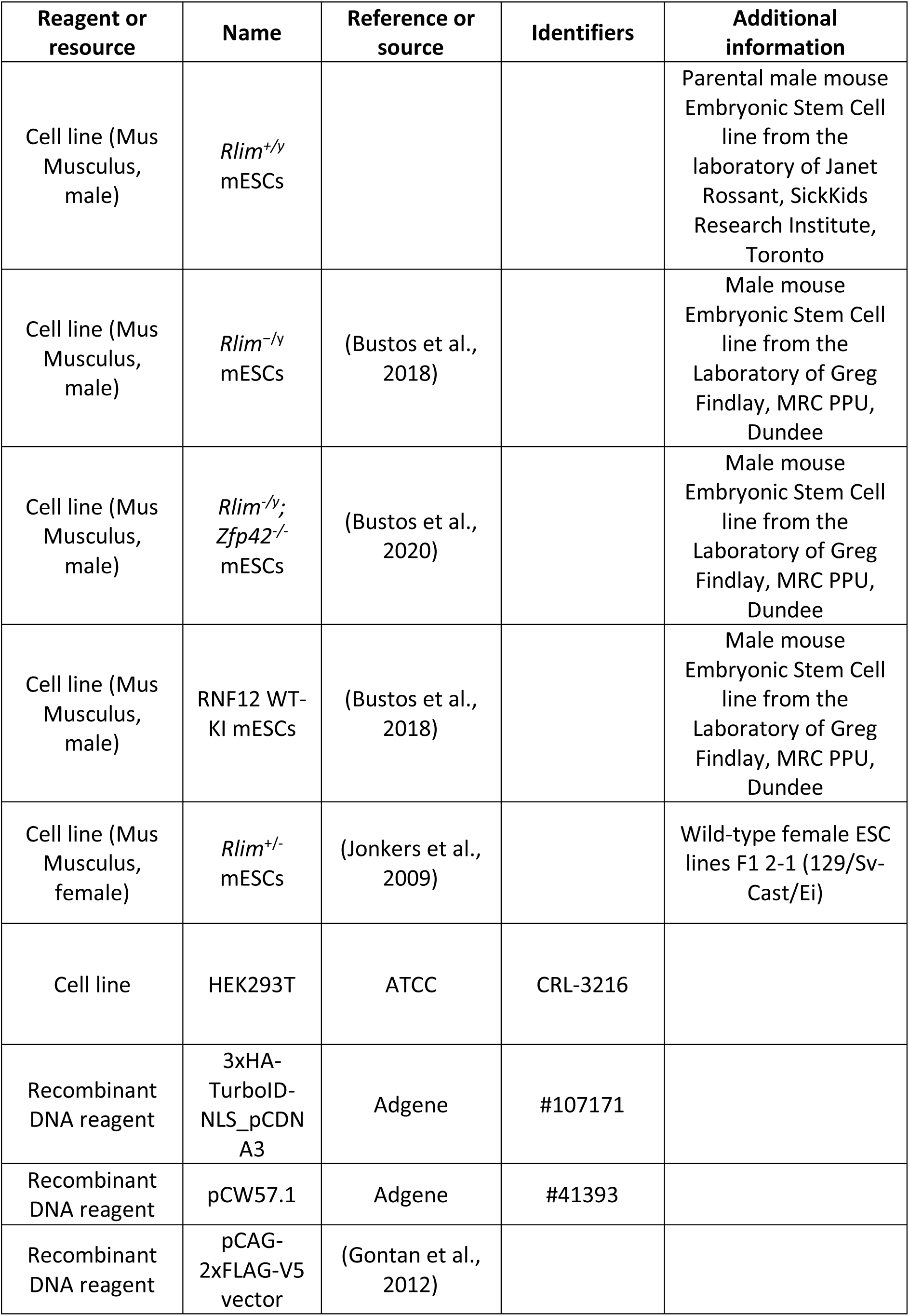
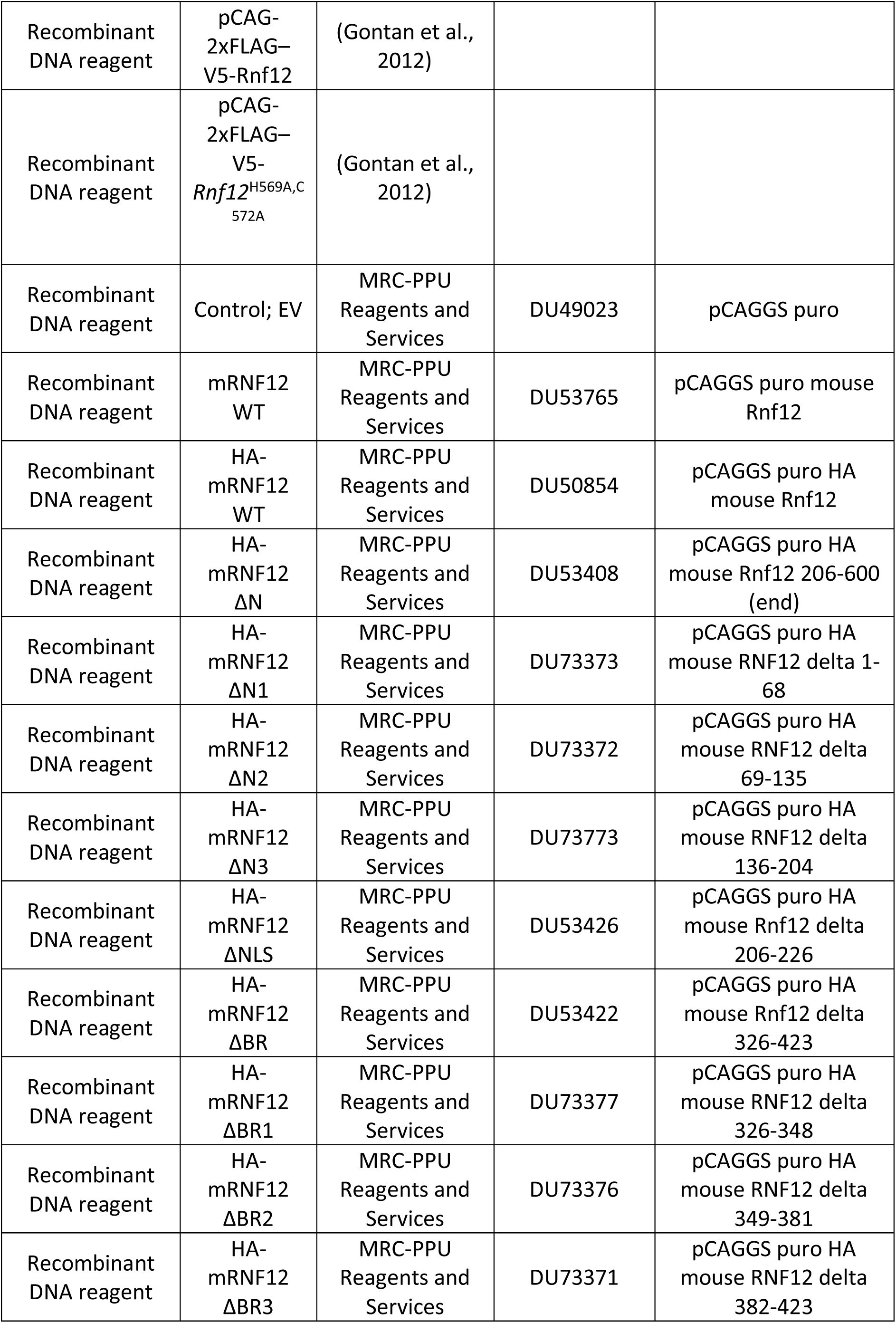
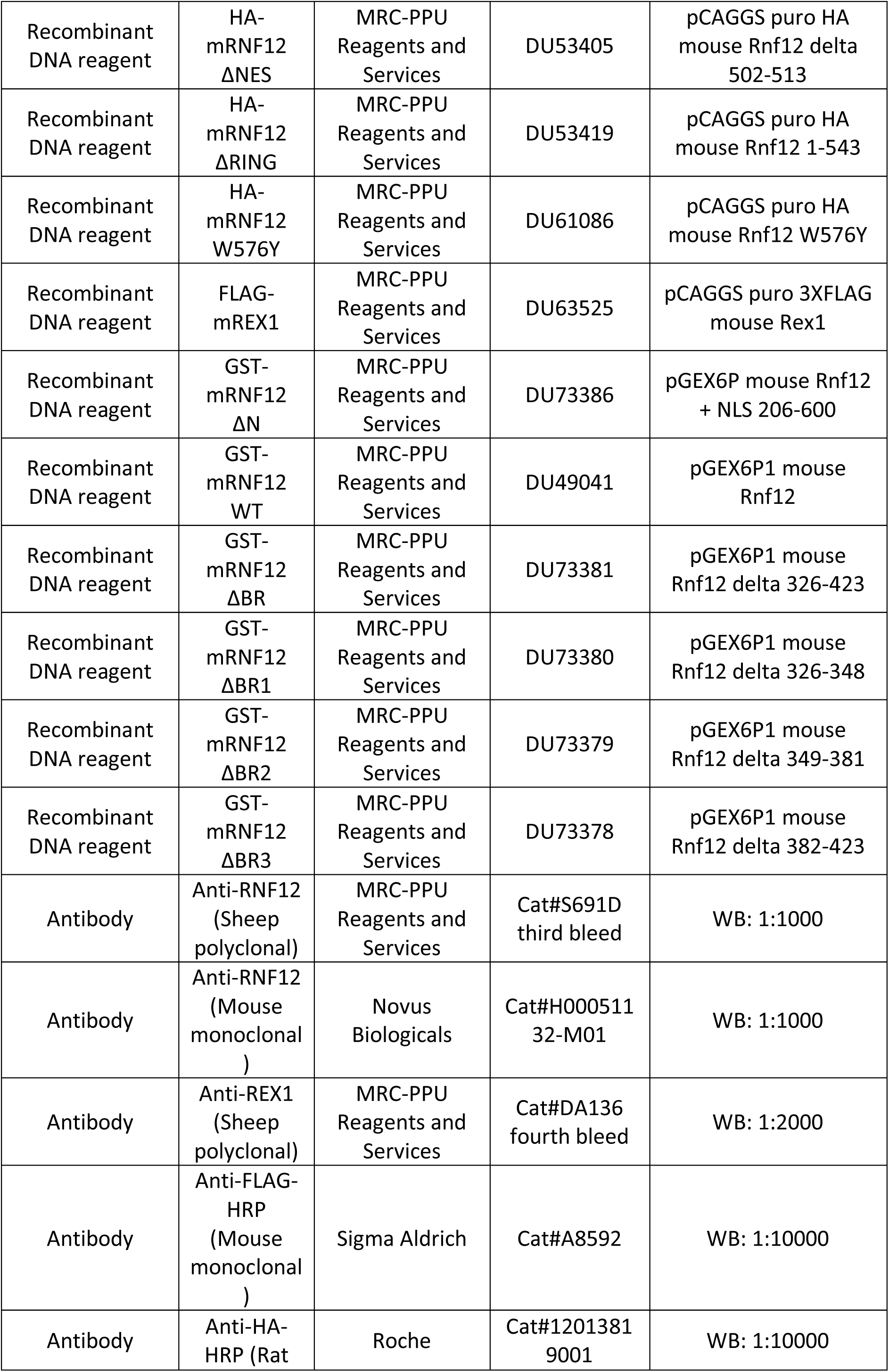
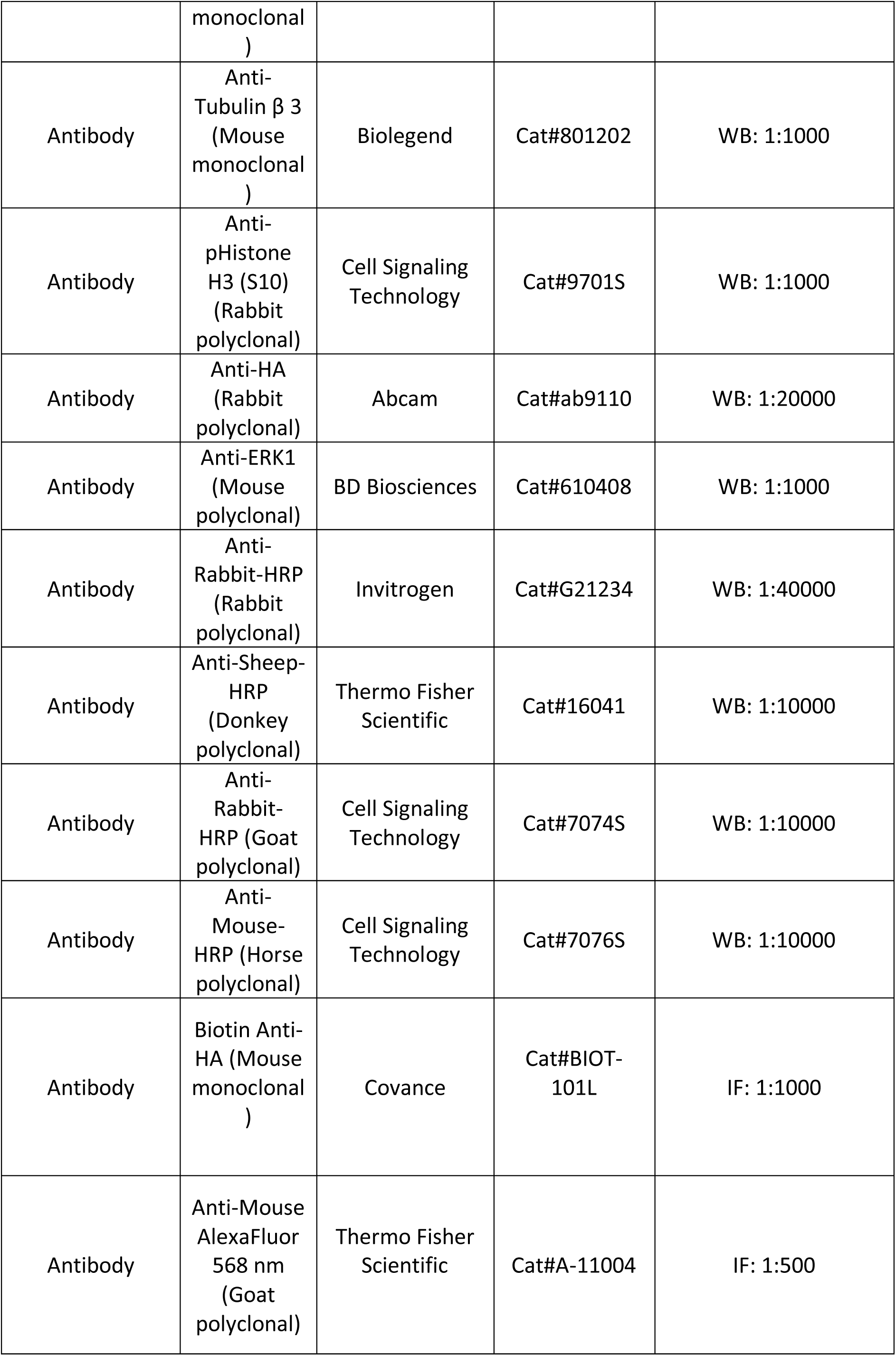
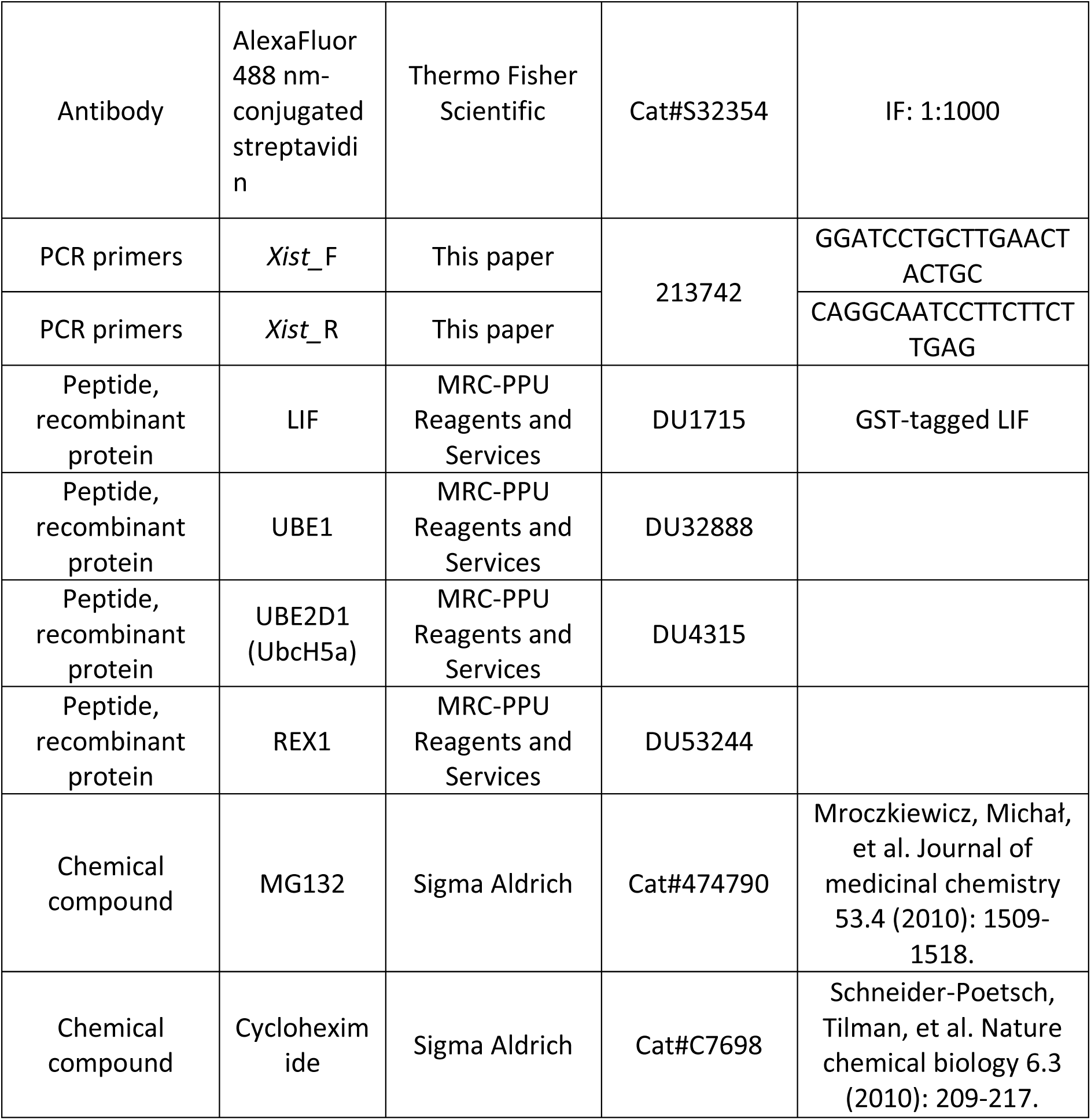
Reagents Summary Table.

### Mouse embryonic stem cell culture and transfection

Male mouse embryonic stem cells (mESCs) were obtained from the laboratory of Janet Rossant, SickKids Research Institute, Toronto. RNF12 wild-type knock-in (WT-KI) (Bustos et al., 2018), RNF12 knock-out (*Rlim*^−/y^) (Bustos et al., 2018) and RNF12 and REX1 double knock-out (*Rlim^-/y^; Zfp42^-/-^*) (Bustos et al., 2020) mESCs were described previously. mESCs were cultured in 0.1% gelatin (w/v)-coated plates in ES-DMEM containing 10% (v/v) fetal bovine serum, 5% (v/v) Knock-Out serum replacement, 2 mM glutamine, 0.1 mM minimum essential media (MEM), Non-essential amino acids, penicillin/streptomycin, 1 mM sodium pyruvate (all from Thermo Fisher Scientific), 0.1 mM β-mercaptoethanol (Sigma-Aldrich) and 20 ng/mL GST-tagged leukemia inhibitory factor (LIF) (Medical Research Council Protein Phosphorylation and Ubiquitin Unit Reagents and Services (MRC-PPU R&S) http://mrcppureagents.dundee.ac.uk) in a controlled atmosphere at 37°C with 5% CO_2_ in a water-saturated incubator. cDNA plasmids clones were transfected in mESCs with Lipofectamine LTX (Thermo Fisher Scientific) according to the manufacturer instructions.

### cDNA plasmids

TurboID plasmids were made using In-Fusion Recombination (Takara Bio USA, Inc.). 3xHA-TurboID was amplified from 3xHA-TurboID-NLS pcDNA3 (Addgene plasmid #107171) and inserted into empty pCW57.1 (Addgene plasmid #41393) using the NheI and BamHI restriction enzyme (RE) sites, with the addition of AgeI RE site built into the 3’ primer. 3xHA-TurboID pCW57.1 was used as the control plasmid. RNF12 WT was amplified via PCR from pCAGGS RNF12 (MRC-PPU R&S) and inserted into 3xHA-TurboID pCW57.1 at AgeI and BamHI RE sites. All other cDNA plasmids are available from MRC-PPU R&S and were verified by DNA sequencing (MRC-PPU DNA Sequencing Service) using DYEnamic ET terminator chemistry (Amersham Biosciences) on Applied Biosystems 3730 automated capillary DNA sequencers.

### Chromatin immunoprecipitation followed by DNA Sequencing (ChIP-SEQ) cell lines

Cell lines stably expressing 2xFLAG-V5-RNF12 and 2xFLAG–V5-RNF12^H569A,C572A^ were generated by electroporation of *Rlim*^+/-^ female mESCs (also termed *Rnf12*^+/-^ mESCs) F1 2-1 (129/Sv-Cast/Ei) (Jonkers et al., 2009), with pCAG-2xFLAG–V5-*Rnf12* or pCAG-2xFLAG–V5-*Rnf12*^H569A,C572A^ vectors followed by puromycin selection. The coding sequence of *Rnf12* was amplified from mouse ESC cDNA and cloned into a TOPO blunt vector (Invitrogen). RNF12^H569A,C572A^ mutant was generated by PCR site-directed mutagenesis. For mammalian expression, the wild-type and mutant *Rnf12* coding sequences were subcloned into pCAG-2xFLAG-V5 vector.

### TurboID cell lines

TurboID stable mESC lines were generated using lentiviral transduction. HEK293T cells (ATCC, CRL-3216) were transfected with each construct and third generation lentiviral packaging plasmids (Cell BioLabs, VPK-206) using Lipofectamine 3000 (Thermo Fisher Scientific) as per the manufacturer’s recommendation. Transfected cells were incubated at 37°C for 6 h, replenished with fresh medium and further incubated at 32°C for 72 h. The culture media was filtered through a 0.45 μm filter, concentrated via ultra-centrifugation (20,000xg and 4°C), resuspended in growth media and added to mESCs along with Polybrene (4 μg/ml; Santa Cruz Biotechnology, Dallas, TX). 96 h after transduction, puromycin (6 μg/ml; Thermo Fisher Scientific) was added to the select transduced cells. Established cell lines were grown in 20 μg/ml puromycin. All cells were tested monthly for mycoplasma contamination.

### TurboID proximity-labelled protein purification

Large-scale TurboID pulldowns were performed in triplicate as described in (May & Roux, 2019) with four 10 cm dishes per sample instead of two. In brief, four 10cm dishes at 80% confluency were incubated with doxycycline (1 mg/ml) (Thermo Fisher Scientific) for 18 h prior to induction of biotinylation with 50 μM biotin (Sigma Aldrich) for 4 h. Cells were rinsed twice with PBS and lysed in 8 M urea 50 mM Tris pH 7.4 containing protease inhibitor (Thermo Fisher Scientific) and DTT, incubated with universal nuclease (Thermo Fisher Scientific), and sonicated to further shear DNA. Lysates were precleared with Gelatin Sepharose 4B beads (GE Healthcare) for 2 h and then incubated with Streptavidin Sepharose High Performance beads (GE Healthcare) for 4 h. Streptavidin beads were washed four times with 8 M urea 50 mM Tris pH 7.4 wash buffer and resuspended in 50 mM ammonium bicarbonate with 1 mM biotin.

To analyse post-pulldown fractions by immunoblot, 10% of the post-pulldown bead fraction was resuspended in SDS–PAGE sample buffer and boiled for 5 mins. Proteins were separated on 4–20% gradient gels (Mini-PROTEAN TGX; Bio-Rad, Hercules, CA) and transferred to nitrocellulose membrane (Bio-Rad). After blocking with 10% (vol/vol) adult bovine serum and 0.2% Triton X-100 in PBS for 30 mins, the membrane was incubated with rabbit anti-HA antibody (Abcam) overnight, washed with PBS and detected using horseradish peroxidase (HRP)–conjugated anti-rabbit (Invitrogen). The signals from antibodies were detected using enhanced chemiluminescence via a Bio-Rad ChemiDoc MP System (Bio-Rad, Hercules, CA). Following detection of HA, the membrane was quenched with 30% H_2_O_2_ for 30 mins. To detect biotinylated proteins, the membrane was incubated with HRP-conjugated streptavidin (1:40,000; ab7403; Abcam) in 0.4% Triton X-100 in PBS for 45 mins.

### Mass spectrometry analysis

Protein samples were reduced, alkylated, and digested on-bead using filter-aided sample preparation (Wiśniewski, Zougman, Nagaraj, & Mann, 2009) with sequencing grade modified porcine trypsin (Promega). Tryptic peptides were separated by reverse phase XSelect CSH C18 2.5 µm resin (Waters) on an in-line 150 x 0.075 mm column using an UltiMate 3000 RSLCnano system (Thermo Fisher Scientific). Peptides were eluted using a 60 mins gradient from 98:2 to 65:35 buffer A:B ratio (Buffer A = 0.1% formic acid, 0.5% acetonitrile, Buffer B = 0.1% formic acid, 99.9% acetonitrile). Eluted peptides were ionized by electrospray (2.4 kV) followed by mass spectrometric analysis on an Orbitrap Fusion Tribrid mass spectrometer (Thermo Fisher Scientific). MS data were acquired using the FTMS analyzer in profile mode at a resolution of 240,000 over a range of 375 to 1500 m/z. Following HCD activation, MS/MS data were acquired using the ion trap analyzer in centroid mode and normal mass range with normalized collision energy of 28-31% depending on charge state and precursor selection range. Proteins were identified by database search using MaxQuant (Max Planck Institute) label-free quantification with a parent ion tolerance of 2.5 ppm and a fragment ion tolerance of 0.5 Da. Scaffold Q+S (Proteome Software) was used to verify MS/MS based peptide and protein identifications. Protein identifications were accepted if they could be established with less than 1.0% false discovery and contained at least 2 identified peptides. Protein probabilities were assigned by the Protein Prophet algorithm (Nesvizhskii, Keller, Kolker, & Aebersold, 2003). Protein interaction candidates were considered if they were identified in all three triplicate runs and enriched at least 3X over both TurboID-only and bait-specific controls.

### ChIP-seq methodology

The ChIP-seq experiments were performed as described (Soler et al., 2011) with minor modifications. For the RNF12 ChIP-seq experiments, 1 × 10^8^ undifferentiated female ESCs expressing V5-tagged RNF12, V5-tagged RNF12^H569A,C572A^ and control wild-type ESCs were cultured without feeders until they reached 80% confluence. Cells were treated for 3 h with proteasome inhibitor (15 μM MG132). All buffers used contained protease inhibitor cocktail tablet (Roche) and 15 μM MG132. The medium was removed, and cells were washed three times with PBS. Then, cells were cross-linked by incubating with PBS containing 2 mM DSG (Thermo Fisher Scientific) for 45 mins at room temperature (RT) on a rotating platform. After the incubation, cells were washed three times with PBS. In the last wash, formaldehyde was added to 1% final concentration and incubated for 10 mins at RT, followed by the addition of glycine to a final concentration of 0.125 M, and cells were incubated for an additional 5 mins at RT to quench the reaction. Cells were washed twice with ice-cold PBS, then scraped and collected in cold PBS. The fixed cell pellets were resuspended in lysis buffer (10 mM Tris-HCl pH 7.5, 1 mM EDTA, 0.5 mM EGTA) and incubated 10 mins on ice. Samples were sonicated on ice using a Sanyo Soniprep 150 sonicator (amplitude 9, 37 cycles of 15 sec on and 30 sec off) to a DNA fragment size in the range of 300–800 nucleotides. The sonicated chromatin samples were centrifuged at 13,000 rpm for 5 mins at 4°C. Chromatin was then diluted to a final volume of 10 ml with dilution buffer (0.01% SDS, 1.1% Triton X-100, 1.2 mM EDTA, 16.7 mM Tris–HCl pH 8, 167 mM NaCl), precleared and immunoprecipitated overnight at 4°C with 60 μl of pre-blocked V5 agarose beads (Sigma) for each ChIP-seq experiment. Beads were washed twice with low salt buffer (0.1% SDS, 1% Triton X100, 2 mM EDTA, 20 mM Tris–HCl pH 8, 150 mM NaCl), followed by 2 washes with high salt buffer (0.1% SDS, 1% Triton X-100, 2 mM EDTA, 20 mM Tris–HCl pH 8, 500 mM NaCl), 2 washes with LiCl buffer (0.25 M LiCl, 1% Np40, 1% sodium deoxycholate, 1 mM EDTA, 10 mM Tris–HCl pH 8), and 2 washes with TE buffer (10 mM Tris–HCl pH 8, 1 mM EDTA). Each wash step was performed for 10 mins at 4°C on a rotating platform. Chromatin was eluted with 500 µL of Elution Buffer (1% SDS; 0.1 M NaHCO3 in H2O). Chromatin was decrosslinked by adding 20µL of 5 M NaCl and incubating at 65°C for 4 h. Then, 10 µL of 0.5 M EDTA, 20 µL of 1 M Tris–HCl pH 6.5, 20 µg of proteinase K were added and incubated at 45°C for 1 h to degrade proteins. DNA was then Phenol–Chloroform extracted and resuspended in 20 µL of H_2_O. The concentration was then measured. Purified ChIP-DNA was prepared for sequencing according to the Illumina protocol and sequenced on a HiSeq 2000 sequencer (Illumina) with a single read for 36 bp and mapped against the reference Mouse_NCBI37.1_AllChromosomes using eland_extended by Illumina pipeline 1.7.0.

### Pharmacological inhibition

Cycloheximide (CHX) was used at a final concentration of 350 µM and MG132 at a final concentration of 10 µM.

### mESC lysate preparation

mESCs were harvested using lysis buffer (20 mM Tris [pH 7.4], 150 mM NaCl, 1 mM EDTA, 1% NP-40 [v/v], 0.5% sodium deoxycholate [w/v], 10 mM β-glycerophosphate, 10 mM sodium pyrophosphate, 1 mM NaF, 2 mM Na_3_VO_4_, and 0.1U/ml Complete Protease Inhibitor Cocktail Tablets (Roche). BCA Protein Assay Kit (Thermo Fisher Scientific) was used to measure protein concentration of lysates obtained according to manufacturer’s instructions. A bovine serum albumin (BSA) protein curve was used as a standard to calculate protein concentration.

### Chromatin fractionation

Method for separation of the soluble and chromatin fractions was based on (Ballabeni et al., 2004). mESCs were harvested by addition of Trypsin-EDTA (Gibco), transferred to a microcentrifuge tube and washed with cold PBS. Cells were centrifuged at 500 g for 5 mins and the resulting pellet was resuspended in CSK buffer (0.5% Triton X-100, 10 mM HEPES pH 7.4, 100 mM NaCl, 300 mM sucrose, 3mM MgCl_2_, 1 mM EGTA, 0.1 U/ml Complete Protease Inhibitor Cocktail Tablets (Roche), Phosphatase inhibitor (20 µl/ml of 50X Phosphatase inhibitor cocktail (5 mM Sodium fluoride (NaF), 1 mM Sodium orthovanadate (Na_3_VO_4_), 1 mM Sodium pyrophosphate, 1 mM β-glycerophosphate)). Samples were incubated on ice for 5 mins, and after centrifugation at 1350 g for 5 mins, the supernatant (soluble fraction) was saved to a new microcentrifuge tube. Pellet was washed 3x (centrifugations are for 3 mins at 1,350 g) with CSK buffer and the final pellet was resuspended with NaCl buffer (0.1% Triton X-100, 50mM TRIS (pH 7.4), 250mM NaCl, 1mM EDTA, 50 mM NaF, protease and phosphatase inhibitor, 2mM MgCl_2_ and Benzonase (1:500; Sigma Aldrich)). Samples were incubated in NaCl buffer on ice for 30 mins with resuspension every 10 mins. Samples were then centrifuged at maximum speed (16,000 g) for 15 mins and the supernatant (chromatin fraction) saved. The chromatin fraction was then centrifuged at 16,000 g for 15 mins to remove any chromatin contamination and supernatant used for further analysis.

### Immunoprecipitation

For HA-tagged protein immunoprecipitation, 10 µl Pierce Anti-HA Magnetic Beads (Thermo Fisher Scientific) were used. Beads were washed three times with lysis buffer and incubated with 1 mg mESC protein lysate overnight at 4°C. Beads were then washed 3x with lysis buffer supplemented with 500 mM NaCl. In each step, beads were separated using a magnetic stand, and the supernatant was discarded. LDS sample buffer was used to elute proteins bound to beads and samples were heated for 5 mins at 95°C.

### Immunoblotting

Commercial NuPAGE™ 4-12% Bis-Tris SDS-PAGE gels (Thermo Fisher Scientific) were used to load denatured protein samples or protein eluates from pulldown experiments. SDS-PAGE gels were then transferred to polyvinylidene fluoride (PVDF) membranes (Merck Millipore) and incubated with primary antibodies diluted in TBS-T (20 mM Tris pH 7.5, 150 mM NaCl supplemented with 0.2% (v/v) Tween-20 (Sigma Aldrich)) containing 5% non-fat milk buffer (w/v) at 4°C overnight. FLAG-HRP and HA-HRP conjugated primaries antibodies were incubated for 1 h at room temperature. Membranes were then washed 3x with TBS-T and incubated with secondary antibody for 1 h at room temperature. Last, membranes were washed 3x with TBS-T and subjected to chemiluminescence detection with Immobilon Western Chemiluminescent HRP substrate (Merck Millipore) using a Gel-Doc XR+ System (Bio-Rad). Images were analysed and quantified using Image Lab software (Bio-Rad).

### Protein purification

Mouse RNF12 WT, RNF12 W576Y and RNF12 ΔN, RNF12 ΔBR, ΔBR1, ΔBR2 and ΔBR3 mutants were cloned into a pGEX6P1 vector. GST-tagged proteins were purified from BL21-CodonPlus (DE3)-RIPL Competent Cells (Agilent, 230280) as follows; colonies from a LB containing Ampicillin (100 μg/ml) plate were transferred into liquid LB media supplemented with Ampicillin (1:1000 dilution) and were left to growth in a 2 liters flask at 37°C until OD600 reached 0.8. 10 μM Isopropyl β-D-thiogalactoside (IPTG) (Sigma) was then added to induce protein expression and cells were incubated at 15°C, 180 rpm overnight. Cells were then harvested at 4,200 rpm for 30 mins at 4°C, and the resulting pellet was resuspended in 40 ml of lysis buffer (50 mM Tris pH 7.5, 150 mM NaCl, 10% glycerol, 1 mM DTT and 2 tablets of Complete Protease Inhibitor Cocktail Tablets (Roche) per 100 ml lysis buffer). Bacterial cells were lysed by 2 mins sonication with 15 sec pulses on/off, and the resulted sample was centrifuged at 18,000 rpm for 25 mins at 4°C. The supernatant was then incubated with Glutathione Sepharose 4B beads (MRC-PPU R&S) for 90 mins on a rotating wheel at 4°C. Samples were then washed 3x with protein buffer (50 mM Tris pH 7.5, 150 mM NaCl, 10% glycerol, 1 mM DTT) and proteins were GST cleaved and eluted from beads using PreScission Protease (MRC-PPU R&S) at 4°C overnight. Supernatant was separated from beads using a Poly-Prep Chromatography Column (BioRad) and concentrated using an Amicon Ultra-15 Centrifugal Filter Unit 10 KDa molecular weight cut-off (Millipore). Final samples were aliquoted and flash-frozen in liquid nitrogen for storage at -80°C. Recombinant ACHE (acetylcholinesterase) protein was produced by Florent Colomb in Dr. Henry McSorley’s laboratory (School of Life Sciences, University of Dundee) as described previously (Vacca et al., 2020).

### RNF12 *in vitro* ubiquitylation assays

RNF12 recombinant protein (140 nM) was incubated with 20 µl ubiquitylation mix containing 0.1 µM UBE1, 0.05 µM UBE2D1 (UBCH5A), 1.5 µg REX1, 2 µM Cy5-Ubiquitin (South Bay Bio), 0.5 mM Tris (2-carboxyethyl) phosphine (TCEP) pH 7.5, 5 mM ATP, 50 mM Tris pH 7.5 and 5 mM MgCl_2_. Reactions were incubated for 30 mins at 30°C, stopped with 2x LDS-Reducing agent mix and heated for 5 mins at 95°C. REX1 recombinant protein and UBE1 and UBE2D1 enzymes were produced by MRC-PPU R&S and purified via standard protocols (http://mrcppureagents.dundee.ac.uk/).

### Electrophoretic Mobility Shift Assay (EMSA)

pGEX6P1 plasmid DNA was linearized by BamHI and NotI restriction enzymes and further purified with the use of a NucleoSpin Gel and PCR Clean-up kit (Thermo Fisher Scientific) according to the manufacturer’s instructions. 0.05 µg of linearized plasmid was incubated with 0.2-0.8 µg of recombinant proteins in a 10 µl sample volume with TE buffer (10 mM Tris pH8.0, 1 mM EDTA) at 37°C for 1 h. Samples were run on a 0.8% agarose gel and analysed using a Chemidoc Imaging System (Bio-Rad).

### Extraction of RNA and quantitative RT-PCR

mESCs transfected with the indicated cDNA plasmids were cultured for 48 h until confluent. For *Xist* induction analysis, *Rlim^+/y^* mESCs were transfected as described and cultured for 72 h in LIF-deficient media prior to lysis. RNA was extracted using an Omega total RNA extraction kit (Omega Biotek) (column-based system) according to the manufacturer’s instructions. The obtained RNA was then converted to cDNA using the iScript cDNA synthesis Kit (Bio-Rad) according to the manufacturer’s instructions. qPCR primers were ordered from Life Technologies and are 20-24 bp with a melting temperature of 58-62°C. Sequences were either acquired from PrimerBank database (https://pga.mgh.harvard.edu/primerbank/) or designed with the use of the Primer3 software. Specificity of each primer was predicted *in silico* with the use of the NCBI Primer-Blast software (https://www.ncbi.nlm.nih.gov/tools/primer-blast/). qPCR was performed using a SsoFast EvaGreen Supermix (Bio-Rad) in 384-well plates and a CFX384 real time PCR system (Bio-Rad). Each sample consisted of 10 µl of a master mix containing 5.5 µl of SYBR Green, 440 nM forward and reverse primers, 1 µl cDNA and nuclease-free water. Relative RNA levels were expressed using the ΔΔC_t_ method and normalized to *Gapdh* expression. Data was analysed in Excel software and plotted making use of GraphPad Prism 9.3.0 software.

### Immunofluorescence

Cells grown on gelatin-coated glass coverslips were fixed in 3% (wt/vol) paraformaldehyde/phosphate-buffered saline (PBS) for 10 mins and permeabilized by 0.4% (wt/vol) Triton X-100/PBS for 15 min. For labeling fusion proteins, a mouse anti-hemagglutinin (HA) antibody was used (Covance). The primary antibody was detected using Alexa Fluor 568–conjugated goat anti-mouse. Alexa Fluor 488–conjugated streptavidin (Thermo Fisher Scientific) was used to detect biotinylated proteins. DNA was detected with Hoechst dye 33342 (Thermo Fisher Scientific). Coverslips were mounted using 10% (wt/vol) Mowiol 4-88 (Polysciences). Epifluorescence images were captured as z-projections using a Nikon A1R confocal microscope and analysed by the NIS-Elements software.

### Data analysis

Data is presented as mean ± SEM of biological replicates, where individual points represent a single biological replicate. Statistical significance was determined by means of paired t-student’s test. GraphPad Prism V9.00 software was used for representation purposes and differences were statistically significant when p<0.05. Immunoblots were quantified using the densitometric analysis in Image Lab software, and data is presented as mean ± SEM of at least three biological replicates. For quantification of REX1 ubiquitylated signal in the chromatin fraction, only the fourth ubiquitylated band (REX1-Ub^4^) was used for quantification purposes. In qPCR experiments, two technical replicates were included per sample and were shown as an average for quantification.

Downstream analysis of TurboID data was performed using Perseus (version 2.0.10.0) and Curtain 2.0 (curtain.proteo.info, developed by Toan Phung, Dario Alessi lab). The MaxQuant output was loaded into Perseus and the data matrix filtered to remove proteins only identified by site, reverse proteins and potential contaminants. Label-free quantification (LFQ) values were transformed by log2, then data was filtered for valid values (at least two valid values in one group). A two-sample t-test was performed and p-values adjusted using the Benjamini-Hochberg multiple hypothesis correction. Data was then exported for inputting into Curtain to generate a volcano plot and proteins with a >2-fold increase were studied. Identification of genes annotated with nuclear localisation and/or function, and gene set enrichment analysis, were performed using the Database for Annotation, Visualisation and Integrated Discovery (DAVID) bioinformatics tool.

### ChIP-SEQ data analysis

The SNPs in the 129/Sv and Cast/Ei lines were downloaded from the Sanger institute (v.5 SNP142) (Keane et al., 2011). These were used as input for SNPsplit v0.3.4 (Felix Krueger & Andrews, 2016) to construct an N-masked reference genome based on mm10 in which all SNPs between 129/Sv and Cast/Ei are masked. The ChIP-seq reads were trimmed and aligned to the N-masked reference genome using Trim Galore v0.6.7 (F Krueger, 2015) and bowtie2 v2.5.0 (D. Kim, Langmead, & Salzberg, 2015), respectively. SNPsplit was then used to assign the reads to either the 129/Sv or Cast/Ei bam file based on the best alignment or to an unassigned bam file if mapping to a region without allele-specific SNPs. The allele-specific and unassigned bam files were merged into a composite bam file using samtools v1.10 (Li et al., 2009).

Peaks were called from the merged bam files using macs2 v2.2.7.1 (Feng, Liu, Qin, Zhang, & Liu, 2012) callpeak with --broad and default settings. For visualization, the tracks were normalized using macs2 bdgcmp with the Poisson P-value as normalization method. Peaks from the different transcription factors were compared using ChIPseeker v1.34.0 (Yu, Wang, & He, 2015). We plotted the overlap between genes with peaks and the peak annotation using the functions vennplot and plotAnnoBar, respectively. Enrichment at the transcription start sites (TSS) was visualized using plotAvgProf and tagHeatmap. For each transcription factor, we searched for overlapping motifs by running bedtools v2.30.0 (Quinlan & Hall, 2010) getfasta to get the sequences of the peaks and meme-chip v5.5.2 (Machanick & Bailey, 2011) from the meme-3 suite (Bailey, Johnson, Grant, & Noble, 2015) using the JASPAR 2018 motif database (Khan et al., 2018).

## ACKNOWLEDGEMENTS

C E-S was funded by a Wellcome Trust 4-year PhD studentship and a core grant to the MRC-PPU (MC_UU_12016). CA was funded by a core grant to the MRC-PPU (MC_UU_12016). GMF was funded by a Wellcome Trust/Royal Society Sir Henry Dale Fellowship (211209/Z/18/Z). BT, JG and CG were funded by a VIDI grant (09150172110079) from the Dutch Research Council (NWO). The Biochemistry Core at Sanford Research and the The IDeA National Resource for Quantitative Proteomics at UAMS are funded by the National Institutes of Health (grants P20 GM103620 and R24GM137786, respectively). The histology and Imaging core at Sanford Research are funded by the National Institutes of Health (grant 1P30GM145398). We thank Toan Phung (laboratory of Prof. Dario Alessi, MRC-PPU) and Frederic Lamoliatte (MRC-PPU) for help with CURTAIN data analysis, Dr Rachel Toth and Dr Thomas Macartney (MRC-PPU R&S) for cloning, Dr Axel Knebel and Dr James Hastie (MRC-PPU R&S) for protein purification, and Florent Colomb and Dr. Henry McSorley (School of Life Sciences, University of Dundee) for help with EMSA assays and recombinant ACHE protein.

## AUTHOR CONTRIBUTIONS

CE-S and GMF conceived the project, prepared figures and wrote the paper. CE-S designed and performed experiments and performed data analysis for Figs. 2-7. CA designed and performed experiments for Figs. 5&6 and performed data analysis for Fig. 1. CG designed and performed experiments for Fig. 2. BFT performed data analysis for Fig. 2. FB Designed and performed experiments for Fig. 1. RJC, KJR and DM performed cloning, viral production, cell line validation, pulldowns and mass spectrometry analysis for Fig. 1. JG, SGM, FB, CG and GMF provided advice, guidance and funding.

